# Profilin 1 maintains cell cycle fidelity to prevent unscheduled genome doubling and polyploidy in cancer

**DOI:** 10.64898/2026.05.12.724607

**Authors:** Federica Scotto di Carlo, Sharon Russo, Simon Gemble, Noemi Vitale, Anne-Sophie Macé, Iris Harmsen, Maria Assunta Borriello, Floris Foijer, Marcello Manfredi, Renata Basto, Fernando Gianfrancesco

## Abstract

Whole-genome doubling (WGD) represents a major route to genome instability and therapeutic resistance in cancer; yet, the mechanisms enabling genome duplication in p53-proficient cells remain poorly understood. Here, we identify Profilin 1 (*PFN1*) loss as a driver of WGD through impaired mitotic entry. Using FUCCI live-cell imaging and single-cell genomic profiling, we show that PFN1-deficient cells bypass mitosis and undergo endoreplication, generating tetraploid cells. Rather than undergoing stable arrest after mitotic bypass, these genome-doubled cells retain proliferative capacity and proceed through aberrant mitotic divisions, thereby amplifying genomic instability. Proteomic analyses reveal coordinated attenuation of late cell-cycle programs, including downregulation of key mitotic regulators such as CDK1, PLK1 and CKS2, consistent with impaired G2/M transition. Despite accumulating polyploidy, PFN1-knockout cells fail to activate an effective p53 tetraploidy checkpoint and display increased nuclear MDM2, promoting cell-cycle arrest evasion and chemotherapeutic resistance. We supported clinical relevance by an orthotopic osteosarcoma xenograft model, in which PFN1-deficient SaOS2 cells showed enhanced metastatic dissemination, and by pan-cancer TCGA analyses confirming a recurrent association between *PFN1* loss and WGD. Together, these findings identify Profilin 1 as a safeguard of cell-cycle fidelity whose loss enables genome doubling, cellular plasticity and therapy tolerance.

## Introduction

Faithful cell division is essential to preserve genome integrity across generations. Eukaryotic cells have evolved complex surveillance mechanisms to ensure that DNA replication occurs only once per cycle and that each daughter cell inherits a complete and balanced genome^1^. Nevertheless, many cancers display whole-genome doubling (WGD), an event in which the entire chromosome set (2n) is duplicated, creating a tetraploid intermediate (4n) that frequently evolves toward aneuploidy and chromosomal instability^2–4^. WGD is estimated to occur in approximately 37% of primary tumors and 56% of metastatic solid tumors^2^. Although WGD can impose a proliferative burden, it also provides a substrate for rapid adaptation by buffering deleterious mutations and promoting therapy resistance^4^.

Under physiological conditions, polyploidization occurs in specialized cell types such as osteoclasts, hepatocytes, megakaryocytes, and trophoblasts, through mechanisms including cell fusion, mitotic slippage, or endoreplication^5^. In contrast, unscheduled genome doubling is normally prevented by cell cycle checkpoints that restrain aberrant cell cycle progression and halt tetraploid cells in G1. The tumor suppressor p53 plays a central role in this surveillance, enforcing cell cycle arrest in response to tetraploidy, centrosome amplification, or replication-associated stress^6–8^. However, large-scale genomic analyses have revealed that a substantial fraction of WGD events arise in tumors retaining wild-type p53, suggesting the existence of alternative mechanisms that allow genome duplication while evading canonical checkpoint control^9^.

Osteosarcoma (OS), the most common malignant bone tumor, represents one of the cancer types with the highest prevalence of WGD, indicating that genome duplication is a prominent feature of its evolutionary trajectory^10^. We previously identified loss-of-function mutations in the *PFN1* gene, encoding Profilin 1, in osteosarcomas arising from Paget’s disease of bone (OS/PDB)^11^, and we also show that *PFN1* expression is frequently reduced in primary OS not associated with PDB. Profilin 1 has been classically characterized as a regulator of cytoskeleton dynamics being an actin-binding protein; however, accumulating evidence indicates that it also contributes to cell cycle regulation through mechanisms that remain incompletely understood. Notably, Profilin 1 overexpression has been reported to induce G1 arrest^12,13^, and more recent studies implicate Profilin 1 in DNA replication control and mitotic progression^14,15^. Consistent with these observations, we recently demonstrated that Profilin 1 depletion perturbs mitotic execution and cytokinesis, leading to chromosome segregation defects and the formation of tetraploid cells through cytokinesis failure^15^. OS/PDB samples harboring *PFN1* loss display highly aberrant karyotypes and complex structural variants, underscoring a link between Profilin 1 deficiency and genome instability. Importantly, sequencing analyses of our OS/PDB cohort revealed that more than half of tumors exhibited WGD events, and in all cases but one, *PFN1* loss occurred prior to genome doubling, suggesting that Profilin 1 inactivation represents an early and potentially permissive event in genome reduplication^15^. However, the extent and dynamics of tetraploidization observed upon *PFN1* depletion cannot be fully explained by cytokinesis failure alone, whose detected frequency was nearly 10% (ref.^15^), raising the possibility that additional cell cycle alterations contribute to genome doubling.

Here, we show that Profilin 1 loss enables WGD through mitotic bypass, with cells exiting G2 into a G1-like state without cell division, and subsequently undergoing endoreplication, thereby re-replicating DNA. This establishes a replication-based route to polyploidization in p53-proficient cells. By integrating live-cell imaging, single-cell genomics, and proteomics analyses, we identify a coordinated alteration of late cell cycle regulatory programs, including impaired G2/M transition and mitotic execution, as a molecular framework underlying this process. Furthermore, we uncover that PFN1-deficient cells evade p53-mediated checkpoint activation through aberrant MDM2 regulation, promoting chemotherapeutic resistance. Together, these findings position Profilin 1 as a critical regulator of cell cycle fidelity whose loss licenses genome doubling and fosters tumor evolution.

## Results

### 1. Profilin 1 loss is associated with whole-genome doubling and drives progressive polyploidization

Whole-genome doubling (WGD) is a prevalent event across human cancers, and the genomic loss of *PFN1* is likewise recurrently observed in multiple tumor types (**Supplementary Fig. 1a,b**). To explore the clinical relevance of Profilin 1 loss in tumor evolution, we analyzed TCGA datasets for correlations between *PFN1* deletion and WGD events across cancer types. Across pan-cancer samples, *PFN1* deletion was significantly enriched in WGD-positive tumors compared to WGD-negative tumors (Fisher’s exact test; OR = 3.33; *p* = 2.0 × 10^-177^) (**Fig. 1a**; **Supplementary Data 1**). Tumor-type–specific analyses further confirmed positive associations in multiple malignancies, including uterine endometrial (OR = 12.99, FDR = 2.9 × 10^-26^), colon (OR = 6.72, FDR = 3.7 × 10^-18^), stomach (OR = 5.49, FDR = 3.7 × 10^-15^), prostate (OR = 7.29, FDR = 2.7 × 10^-7^), breast (OR = 1.99, FDR = 6.9 × 10^-7^), and glioblastoma (OR = 4.13, FDR = 2.8 × 10^-6^) cancers (**Fig. 1a**; **Table 1**; **Supplementary Fig. 1c**). Collectively, these findings indicate a robust and recurrent association between *PFN1* genomic loss and WGD across diverse tumor contexts.

**Figure 1:**
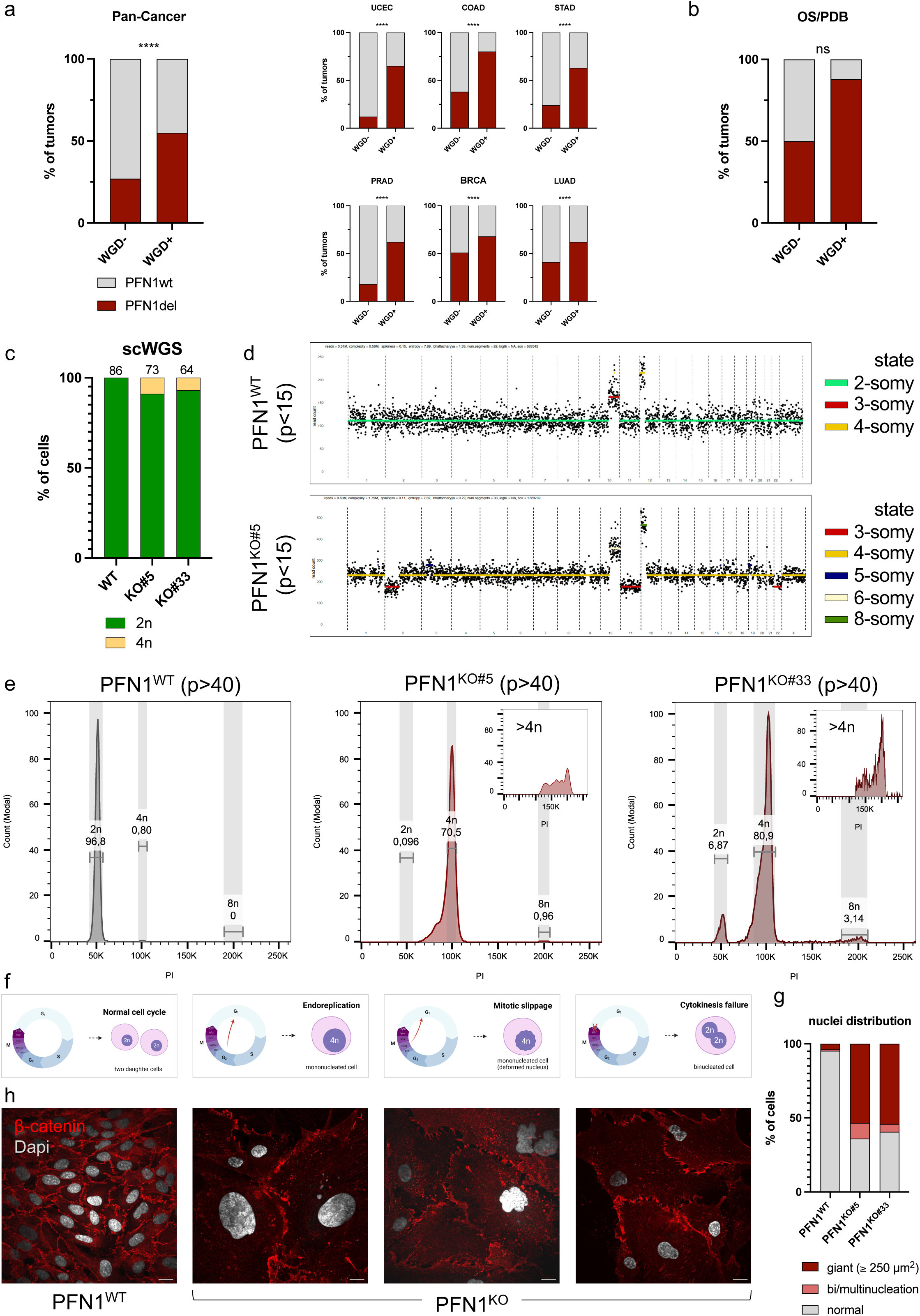
Profilin 1 loss is associated with whole-genome doubling and progressive changes in cellular ploidy **a.** Stacked bar plots showing the distribution of tumors with or without whole-genome doubling (WGD), stratified by PFN1 status (WT or DEL) across pan-cancer samples (**left**) and selected cancer types (1a, **right**). Data are expressed as the percentage of tumors within each WGD category. Cancer types are indicated as follows: Uterine Corpus Endometrial Carcinoma (UCEC), Colon Adenocarcinoma (COAD), Stomach Adenocarcinoma (STAD), Prostate Adenocarcinoma (PRAD), Breast Invasive Carcinoma (BRCA), and Lung Adenocarcinoma (LUAD). Asterisks indicate statistical significance assessed by two-sided Fisher’s exact test (*p* < 0.0001). *P*-values were calculated using absolute tumor counts, whereas percentages are shown for visualization. Corresponding odds ratios (OR), 95% confidence intervals (CI), and false discovery rate (FDR)–adjusted *p*-values are reported in **Table 1**. **b.** Stacked bar plots showing the distribution of OS/PDB tumors with or without WGD stratified by PFN1 status (WT or DEL). Data are expressed as the percentage of tumors within each WGD category. Statistical significance was assessed using a two-sided Fisher’s exact test (*p* = 0.245) calculated using absolute tumor counts, whereas percentages are shown for visualization; sample size and statistical details are provided in **Supplementary Data 1**. **c.** Stacked bar plot representing the distribution of ploidy states inferred from scWGS in early-passage (p<15) serum-starved PFN1^WT^ and two independent PFN1^KO^ RPE1 clones (#5, #33). Bars indicate the relative proportion (%) of diploid (2n; green) and tetraploid (4n; yellow) cells. Total number of single cells sequenced is shown on each bar. **d.** Representative single-cell whole-genome copy-number profiles from serum-starved RPE1 cells, showing a diploid (2n) PFN1^WT^ cell (top) and a tetraploid (4n) PFN1^KO#5^ cell (bottom). Data points are color-coded according to inferred ploidy states; different ploidy states are indicated by distinct colors. **e.** DNA content analysis by propidium iodide (PI) staining in serum-starved late-passage (p>40) WT and PFN1^KO^ RPE1 cells. Histograms show cell counts expressed in modal values plotted against DNA content measured by PI incorporation. Diploid (2n), tetraploid (4n), and octoploid (8n) peaks are highlighted and relative abundance (%) is shown. Insets emphasize the presence of heterogeneous DNA content exceeding discrete ploidy states. **f.** Schematic overview of potential cellular routes leading to polyploidization, including endoreplication, mitotic bypass, and cytokinesis failure. **g.** Quantification of cells with normal (< 250 µm^2^), giant (β 250 µm^2^), or multiple nuclei in PFN1^WT^ and two PFN1^KO^ RPE1 cell populations, expressed as percentage of total cells. *N* = 635 cells for WT; 492 cells for KO#5; 532 cells for KO#33 across at least three independent experiments. **h.** Representative images of WT and PFN1^KO^ cells stained with DAPI (grey) and β-catenin (red) to delineate cell borders, illustrating differences in nuclear size and morphology. Scale bars, 20 µm.

**Table 1:** Association between WGD and *PFN1* loss across tumor types.

| Tumor | Total_N | OR | 95% CI | FDR |
| --- | --- | --- | --- | --- |
| UCEC | 531 | 12.99 | 8.02–21.03 | 2.9e-26 |
| COAD | 445 | 6.72 | 4.31–10.49 | 3.7e-18 |
| STAD | 429 | 5.49 | 3.60–8.37 | 3.7e-15 |
| PRAD | 472 | 7.29 | 3.59–14.79 | 2.7e-07 |
| BRCA | 1049 | 1.99 | 1.54–2.56 | 6.9e-07 |
| GBM | 561 | 4.13 | 2.43–7.03 | 2.8e-06 |
| KIRC | 501 | 4.72 | 2.51–8.86 | 1.4e-05 |
| LUAD | 506 | 2.34 | 1.63–3.35 | 1.6e-05 |
| DLBC* | 47 | 93.33 | 8.55–1018.59 | 1.6e-05 |
| BLCA | 401 | 2.47 | 1.63–3.72 | 5.6e-05 |
| THCA* | 464 | 37.33 | 7.27–191.68 | 0.001 |
| HNSC | 515 | 2.07 | 1.37–3.11 | 0.002 |
| THYM | 106 | 7.96 | 2.02–31.37 | 0.014 |
| PAAD | 165 | 2.78 | 1.30–5.97 | 0.025 |
| CESC | 294 | 1.88 | 1.14–3.10 | 0.032 |
| SKCM | 357 | 1.74 | 1.12–2.71 | 0.035 |

Odds ratios (OR) were calculated using Fisher's exact test based on 2x2 contingency tables (WGD status x *PFN1* deletion). 95% confidence intervals (95% CI) were computed on the log(OR) scale (Woolf method). P-values were adjusted for multiple testing using the Benjamini–Hochberg false discovery rate (FDR). OR > 1 indicates enrichment of *PFN1* deletion in WGD-positive tumors.
\*Tumor types marked with an asterisk had limited sample size and/or low numbers of *PFN1*-deleted cases, which may yield unstable OR estimates (wide 95% CI).
Tumor abbreviations are provided in **Supplementary Data 1**.

To extend these observations to osteosarcoma, we reanalyzed whole-genome sequencing data from our previously published OS/PDB cohort^15^, specifically assessing the relationship between *PFN1* deletion and WGD status in this tumor context. Consistent with the pan-cancer analysis, *PFN1* loss was observed in 87.5% (7/8) of WGD-positive tumors compared to 50% (3/6) of WGD-negative tumors (OR = 7.0; **Fig. 1b**). Despite the small cohort size typical of this rare tumor subtype, the magnitude and direction of the association are consistent with those observed across TCGA cancers, supporting a biologically meaningful link between *PFN1* loss and genome doubling in OS/PDB.

To model this phenomenon experimentally, we monitored DNA content in hTERT-RPE1 (RPE1) cells previously characterized^15^ for *PFN1* knockout (PFN1^KO^) at different passages using propidium iodide (PI) staining. In serum-starved early-passage (p<15) PFN1^KO^ cells, the DNA profile resembled wild-type (WT) cells, with a predominantly diploid population and only a minor fraction (< 5%) of 4n cells detected consistently across both independent clones (**Supplementary Fig. 2a**). Because serum starvation arrests RPE1 cells predominantly in G0/G1, diploid cells are expected to display a 2n DNA content under these conditions. Therefore, the detection of 4n cells in serum-starved cultures is unlikely to reflect a transient diploid G2/M population and instead indicates the presence of tetraploid cells. This early appearance of a small 4n population suggests the initial emergence of tetraploid cells soon after *PFN1* loss. To directly assess DNA copy number at single-cell resolution and confirm the presence of tetraploid cells, we performed single-cell whole-genome sequencing (scWGS) on serum-starved early-passage (p<15) PFN1^KO^ cells (two independent clones) compared to WT RPE1 cells (**Supplementary Fig. 2b,c**). Among 137 analyzed KO single cells, 7.3% (10 cells) exhibited tetraploidy (4n), consistent with flow-cytometry analysis (**Fig. 1c**). Despite their nominally tetraploid DNA content, individual cells displayed widespread chromosomal imbalances and structural variations, including random gains and losses (3n–8n), indicative of chromosomal instability even at early stages (**Fig. 1d; Supplementary Fig. 2d**). These alterations occurred in addition to the pre-existing gains of chromosome 10q and chromosome 12 in the parental RPE1 cells^16^. To assess how tetraploidization evolves over time, we analyzed serum-starved PFN1^KO^ clones by PI staining at later passages, compared to matched WT cells (p>40). These cells displayed a marked shift toward higher DNA content, with the majority of cells (70-80%) showing 4n and > 4n peaks (**Fig. 1e**). Notably, inspection of DNA content profiles revealed the presence of cell populations with heterogeneous DNA content exceeding 4n, indicating that PFN1-deficient cells do not adopt stable tetraploid states but instead undergo progressive chromosomal gains and losses, consistent with ongoing genomic instability (**Fig. 1e, insets**). This gradual enrichment of polyploid cells demonstrates that *PFN1* loss drives progressive polyploidization.

To delineate the potential mechanisms underlying this process, we considered different routes to tetraploidization, schematized in **Fig. 1f**: endoreplication, mitotic slippage, and cytokinesis failure. These mechanisms can be morphologically distinguished, as endoreplication results in mononucleated cells with enlarged, round nuclei^17^; mitotic slippage leads to cells with large, often irregular nuclei^18^; and cytokinesis failure typically produces binucleated cells that persist for one cell cycle^19^. Quantification of binucleated cells in PFN1^KO^ cultures revealed that cytokinesis failure accounts for fewer than 10% of events, consistent with the frequency observed by live-cell imaging in our previous study^15^ (**Fig. 1g**), suggesting that additional mechanisms contribute to genome doubling. Indeed, morphological inspection of DAPI-stained cultures revealed a prevalence of giant, mononucleated cells (β 250 µm^2^), as opposed to bi- or multinucleated cells, further supporting that polyploidization primarily arises through endoreplication or mitotic slippage rather than cytokinetic defects (**Fig. 1g,h; Supplementary Fig. 2e**).

Collectively, these findings demonstrate that Profilin 1 loss correlates with WGD in human tumors and drives the emergence of tetraploid cells that progressively evolve toward heterogeneous polyploid states in vitro, establishing a mechanistic link between Profilin 1 deficiency, genome doubling, and chromosomal instability.

### 2. Profilin 1 depletion induces genome duplication through mitotic bypass

To dissect how Profilin 1 loss leads to tetraploidization, we employed the fluorescent ubiquitination-based cell cycle indicator (FUCCI(CA)2) system, which labels G1 (Cdt1^high^, red), S (Geminin^high^, green), and G2 (Cdt1^high^/Geminin^high^, yellow) phases, enabling real-time monitoring of cell cycle dynamics in individual cells (**Fig. 2a; ref.**^20^). Long-term live-cell imaging of PFN1^WT^ and PFN1^KO^ RPE1 FUCCI cells was performed at early/mid passages (p=20-30) to capture the cellular events that ultimately lead to the full tetraploidization observed during clonal evolution (p>40). Our analysis revealed striking differences in cell cycle progression between the two genotypes. WT RPE1 cells cycled regularly through G1–S–G2–M–G1, with an average cell cycle duration of ∼21 hours (**Fig. 2b, top panels**; **Fig. 2c, left**; **Supplementary Video 1**). During long-term live imaging, WT cells frequently completed multiple consecutive divisions, allowing the tracking of up to three successive generations within the same lineage (**Fig. 2c**). Notably, daughter cells originating from the same division typically displayed highly symmetric cell cycle behavior, progressing through subsequent phases with comparable timing and fate (**Fig. 2c**). In contrast, PFN1-deficient cells displayed profound alterations in cell cycle execution. Approximately one-third of PFN1^KO^ cells underwent mitotic bypass (MB), a phenotype characterized by Geminin degradation in the absence of detectable mitotic entry: in these cells, no evidence of nuclear envelope breakdown was observed, as evidenced by the lack of cytoplasmic dispersion or leakage of the FUCCI fluorescent reporters, suggesting a direct transition from G2 into a G1-like state without chromosome segregation (**Fig. 2b, bottom panels**; **Fig. 2c, right**; **Supplementary Fig. 3a**). Importantly, MB was not associated with a single uniform outcome but instead gave rise to distinct cell-fate trajectories during the ∼72-hour imaging window. In some cases, after the mitotic bypass, cells remained in G1 for the remaining observation window (**Fig. 2c; Supplementary Video 2**), whereas in others, mitotic bypass was followed by re-entry into S phase, completing an endoreplication cycle (**Fig. 2c; Supplementary Video 3**). Notably, a subset of cells that had bypassed mitosis and undergone endoreplication subsequently progressed through an apparently complete division in the following cycle (**Fig. 2c; Supplementary Video 4**), indicating that mitotic bypass does not irreversibly commit cells to cell cycle arrest or terminal endocycling. Consistent with the heterogeneous nature of this response, daughter cells originating from the same division occasionally adopted divergent fates, with one cell undergoing MB while its sister cell continued cycling or entered cell cycle arrest (**Supplementary Video 5**). In rare cases, lineage-tracing further revealed that cytokinesis failure and mitotic bypass can occur sequentially within the same cell lineage **(Supplementary Video 6**). This asymmetric behavior underscores cell-to-cell variability in response to *PFN1* loss, even within a shared lineage. Importantly, because MB occurs after completion of a prior S phase, cells exiting G2 without mitosis inevitably enter G1 with a tetraploid DNA content, establishing mitotic bypass as a direct route to genome doubling.

**Figure 2:**
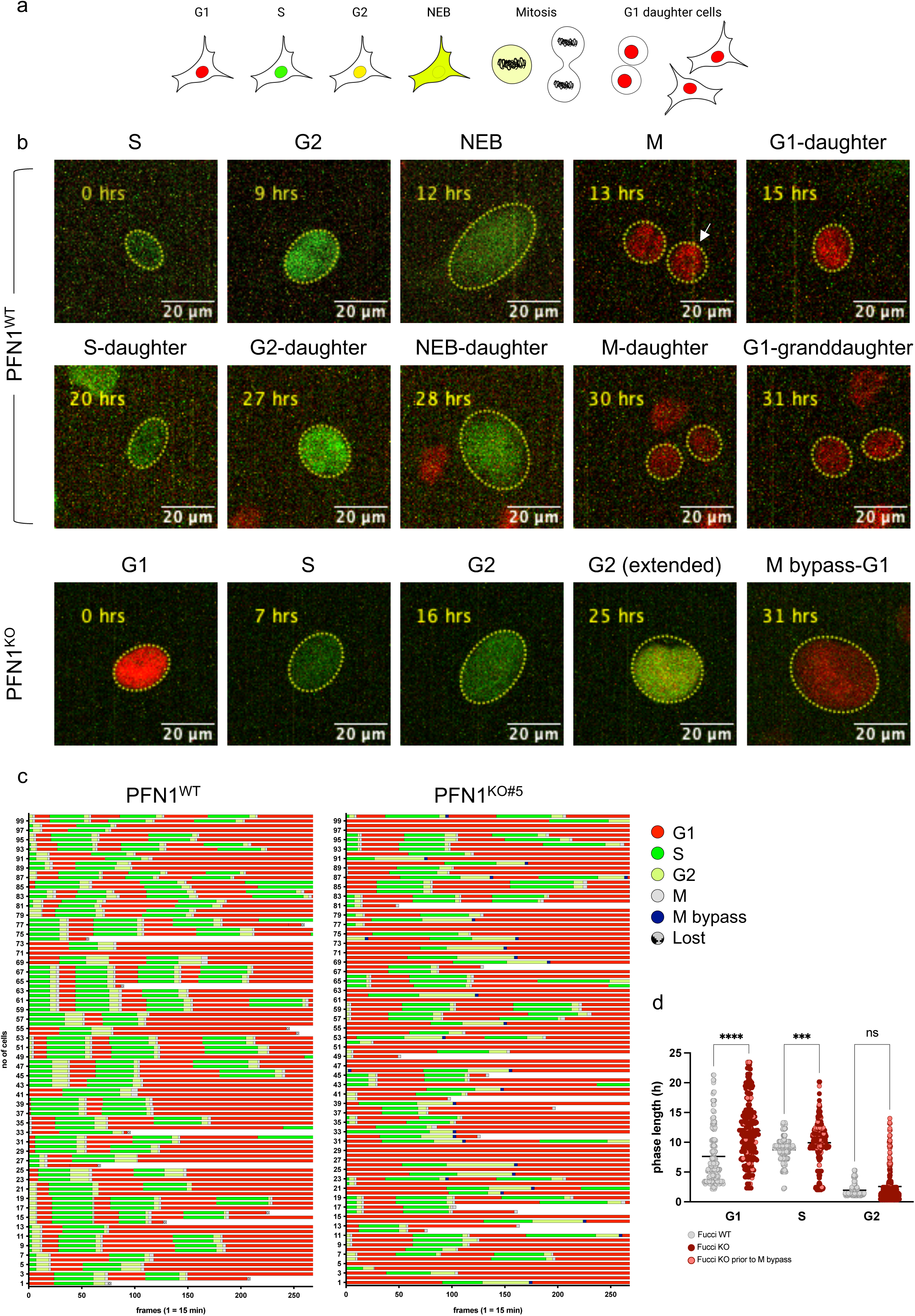
PFN1 deletion drives mitotic bypass **a.** Schematic overview of the FUCCI(CA)2 cell cycle reporter system, illustrating the fluorescent signals associated with the different cell cycle phases. Red = G1; green = S; yellow = G2; dispersion of fluorescence = Nuclear Envelope Breakdown (NEB). **b.** Representative still images from long-term time-lapse imaging of FUCCI-expressing RPE1 cells. Top panels show WT cells, and bottom panels show PFN1^KO^ cells, illustrating cell cycle dynamics over time. Yellow dashed outlines indicate the tracked nucleus, or the corresponding cell during nuclear envelope breakdown (NEB) when FUCCI fluorescence is no longer confined to the nucleus. Time is indicated in hours and was assigned at the onset of fluorescence color transition, corresponding to entry into the subsequent cell cycle phase. Scale bar, 20 µm. **c.** Duration and sequence of cell cycle phases in WT (left) and PFN1^KO#5^ (right) RPE1 cells, as determined by FUCCI-based tracking. Each horizontal lane represents a single cell (*n* = 100 cells per condition). The x-axis represents time in imaging frames (1 frame = 15 minutes). Cell cycle phases are color-coded as follows: G1 (red), S (green), G2 (yellow), mitosis (grey), mitotic bypass (blue), and lost cells (grey with black dots). **d.** Dot plots showing the duration (in hours) of individual cell cycle phases in PFN1^WT^ (grey) and PFN1^KO^ (dark red) RPE1 cells. PFN1^KO^ cells that subsequently undergo mitotic bypass are highlighted in pink. For PFN1^KO^, data from KO#5 and KO#33 clones were pooled. The number of analyzed cells was as follows: WT, *n* = 166 (G1), 179 (S), and 177 (G2); KO, *n* = 290 (G1), 316 (S), and 364 (G2). Statistical significance was assessed using an ordinary one-way ANOVA; adjusted *p*-values are as follow: G1 *vs* G1, *p* < 0.0001; S *vs* S, *p* = 0.0009; G2 *vs* G2, *p* = 0.1751.

Quantitative analysis of FUCCI traces revealed that PFN1^KO^ cells exhibited an overall lengthening of the cell cycle compared to WT cells, with a selective and pronounced extension of the G2 phase specifically in cells undergoing MB (**Fig. 2d**; **Supplementary Fig. 3b**). This selective delay suggests an impaired G2/M transition in PFN1-deficient cells, whereby cells attempt to engage mitotic entry but ultimately exit G2 without chromosome segregation.

### 3. Profilin 1 loss drives mitotic bypass–dependent endoreplication and aberrant cell cycle re-entry

To evaluate whether mitotic bypass was coupled to DNA synthesis and to validate that the Geminin-dependent fluorescence detected by the FUCCI system accurately reflects S-phase progression, we first quantified nuclear size in G1-phase cells before and after bypass events in the same individual cells tracked over time. PFN1^KO^ cells displayed a consistent increase in nuclear area following MB, with an average ∼2.2-fold enlargement compared to pre-bypass G1 nuclei (**Fig. 3a,b**). This increase supports the conclusion that mitotic bypass is not merely associated with cell cycle exit but is accompanied by a *bona fide* increase in DNA content, reflected by nuclear expansion. Notably, analysis of nuclear size in G1-phase cells prior to MB revealed substantial heterogeneity within the PFN1^KO^ population. Sixty-two percent of cells exhibited G1 nuclear areas below 250 µm², consistent with cells undergoing MB for the first time (**Fig. 3b,c**). In contrast, the remaining 38% of cells displayed markedly enlarged G1 nuclei (≥ 250 µm²) already before the observed MB, suggesting that these cells had already undergone at least one prior round of MB and endoreplication before the period of observation (**Fig. 3c; Supplementary Video 7**).

**Figure 3:**
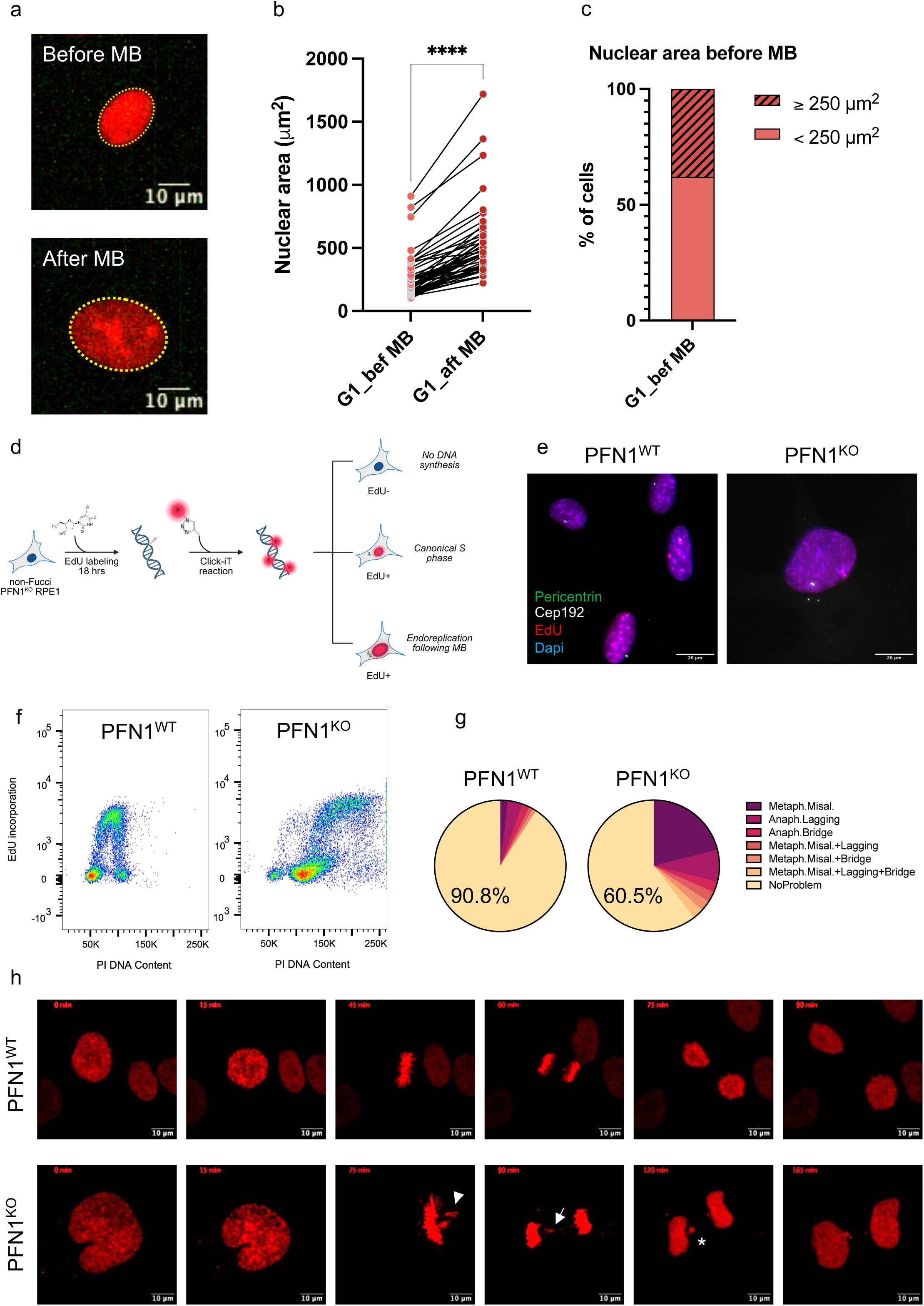
Profilin 1 depletion licenses endoreplication and aberrant proliferative cycling **a.** Representative still images of a PFN1^KO^ RPE1 cell in G1 phase before (top) and after (bottom) a mitotic bypass event, illustrating the increase in nuclear size within the same cell tracked over time. Nuclear contours are outlined in yellow. Scale bar, 10 µm. **b.** Quantification of G1 nuclear area in PFN1^KO^ RPE1 cells before and after mitotic bypass (MB), measured longitudinally in the same individual cells. Statistical significance was assessed using a paired t-test (*n* = 50 cells); *p* < 0.0001. **c.** Stacked bar plot showing the frequency of cells with normal-sized nuclei (< 250 µm²) and giant-sized nuclei (≥ 250 µm²) prior to the filmed mitotic bypass. **d.** Schematic overview of EdU staining and data interpretation criteria. EdU-negative (EdU⁻) cells did not undergo DNA synthesis; EdU-positive (EdU⁺) cells with normal-sized nuclei (< 250 µm²) and ≤ 2 centrosomes underwent canonical S phase; EdU⁺ cells with giant-sized nuclei (≥ 250 µm²) and ≥ 3 centrosomes underwent mitotic bypass followed by endoreplication. **e.** Representative images of EdU⁺ PFN1^WT^ and PFN1^KO^ RPE1 cells. Centrosomes are identified by co-staining for Cep192 (white) and Pericentrin (green). Nuclei are counterstained with DAPI (blue). Scale bar, 20 µm. **f.** Flow cytometry analysis of EdU incorporation combined with DNA content measurements in late-passage PFN1^WT^ and PFN1^KO#33^ RPE1 cells. Each plot displays EdU fluorescence versus propidium iodide (PI) signal, enabling the visualization of DNA synthesis in cells with different DNA contents. **g.** Quantification of mitotic defects identified by time-lapse microscopy in PFN1^WT^ and PFN1^KO^ RPE1 cells expressing H2B–mCherry. Mitotic errors were classified as metaphase misalignment, anaphase lagging chromosomes, anaphase bridges, or combinations thereof. Percentages refer to the frequency of mitoses without detectable chromosome segregation errors. A total of 109 WT and 114 KO cells were analyzed. **h.** Representative still images of H2B–mCherry–expressing PFN1^WT^ (top) and PFN1^KO^ (bottom) RPE1 cells during mitosis. Representative examples of metaphase misalignment, chromatin bridge formation, and micronucleus formation are shown, indicated by an arrowhead, arrow, and asterisk, respectively. Time is indicated in minutes. Scale bar, 10 µm.

To further confirm that mitotic bypass was accompanied by actual DNA replication, we performed EdU incorporation assays in non-FUCCI PFN1^WT^ and PFN1^KO^ cells following an 18-hour labeling period. In this setting, enlarged nuclear size and the presence of supernumerary centrosomes were used as morphological proxies of prior mitotic bypass and DNA re-replication (**Fig. 3d**). A substantial fraction of PFN1^KO^ cells meeting these criteria incorporated EdU, indicating re-entry into S phase (active DNA synthesis) following MB (**Fig. 3e**; **Supplementary Fig. 3c**). As an additional control, cells were co-stained for PCNA, which showed a nuclear pattern consistent with active DNA replication^21^ (**Supplementary Fig. 3d**). Quantitative analysis of EdU fluorescence intensity further showed that, although PFN1^KO^ cells incorporated EdU, the overall signal was shifted toward lower intensity ranges compared to WT cells (**Supplementary Fig. 3c**), suggesting reduced DNA synthesis per replication round.

To further validate that DNA replication occurred in cells with already replicated genomes, we also performed flow cytometry analysis of EdU incorporation in combination with DNA content measurements. In PFN1^WT^ cells, EdU-positive events were associated to canonical S-phase progression within a diploid cell cycle. In contrast, PFN1^KO^ cells displayed a distinct population of EdU-positive cells with a tetraploid DNA content (**Fig. 3f**; **Supplementary Fig. 3e**). This pattern indicates that a substantial fraction of PFN1-deficient cells continues to undergo DNA synthesis despite having already duplicated their genome, consistent with an endoreplication phenotype.

Finally, because the FUCCI system does not allow direct visualization of chromosome condensation and segregation dynamics, we complemented these analyses by performing live-cell imaging of WT and PFN1^KO^ cells expressing H2B-mCherry to assess mitotic behavior and chromosomal integrity during division, particularly in tetraploid/polyploid cells. Using this approach, PFN1^KO^ cells with enlarged nuclei were observed to re-enter mitosis and attempt cell division; however, these divisions were frequently aberrant, displaying chromosome segregation defects such as mis-alignment on the metaphase plate and lagging chromosomes (**Fig. 3g,h; Supplementary Videos 8,9**).

Together, these findings demonstrate that Profilin 1 knockout enables p53-proficient cells to bypass mitosis and reinitiate DNA replication, generating a population of polyploid, genomically unstable cells that retain proliferative capacity despite abnormal karyotypes. Importantly, mitotic bypass in PFN1-deficient cells does not result in terminal arrest: cells can re-enter S phase and attempt subsequent mitotic divisions, converting mitotic bypass from a safe arrest mechanism into a source of genome reduplication and chromosomal instability.

### 4. Profilin 1 loss impairs p53 checkpoint activation through MDM2 upregulation and confers chemotherapeutic resistance

Cells that acquire abnormal genome content typically undergo a p53-mediated cell cycle arrest, commonly triggered by features such as centrosome amplification and increased cell size, as observed in enlarged cells following CDK4/6 inhibition^22–25^. We therefore asked whether PFN1-deficient cells retain this canonical p53-dependent response to abnormal cellular states. We analyzed p53 levels in relation to nuclear size and centrosome number, using these features as a proxy for endoreplicated cells. Immunofluorescence analysis revealed that, although p53 was detectable in PFN1^KO^ cells, p53 levels did not rise substantially, even in cells displaying increasing nuclear size or centrosome amplification (**Fig. 4a,b**), indicating an attenuated and uncoupled p53 response in tetraploid and endoreplicating cells. Consistent with this observation, the E3 ubiquitin ligase MDM2, a key negative regulator of p53 stability, was upregulated in PFN1^KO^ cells and displayed prominent nuclear localization (**Fig. 4c,d; Supplementary Fig. 4a**), suggesting sustained p53 suppression even in the presence of abnormal genome content. To further evaluate p53 responsiveness, WT and PFN1^KO^ cells were exposed to Nutlin-3a (an MDM2 inhibitor), Doxorubicin (Doxo, a DNA-damaging agent that inhibits topoisomerase II and induces DNA breaks), or Hydroxyurea (HU, which activates the DNA damage checkpoint by depleting dNTP pools). Western blot analysis showed that p53 accumulation was markedly reduced in PFN1^KO^ cells compared with WT across all treatments (**Fig. 4e**; **Supplementary Fig. 4b,c**). Consistent with impaired p53 activation, transcriptional targets of p53, including MDM2 and p21, were also poorly induced upon Nutlin-3a treatment (**Fig. 4e**), indicating a blunted p53 transcriptional response and excluding the possibility that the elevated basal MDM2 levels observed in PFN1^KO^ cells (**Fig. 4c,d**) are driven by p53 activity. Similarly, γH2AX induction and phosphorylation of CHK1 and CHK2 were attenuated in PFN1^KO^ cells following HU and Doxo treatment, suggesting a weakened DNA damage response (**Supplementary Fig. 4c**). Together, these findings indicate that although the p53 pathway is not completely inactive in PFN1-deficient cells, its activation is attenuated and not specifically dependent on polyploidization and replication-associated stress, potentially allowing continued proliferation despite abnormal genome content.

**Figure 4:**
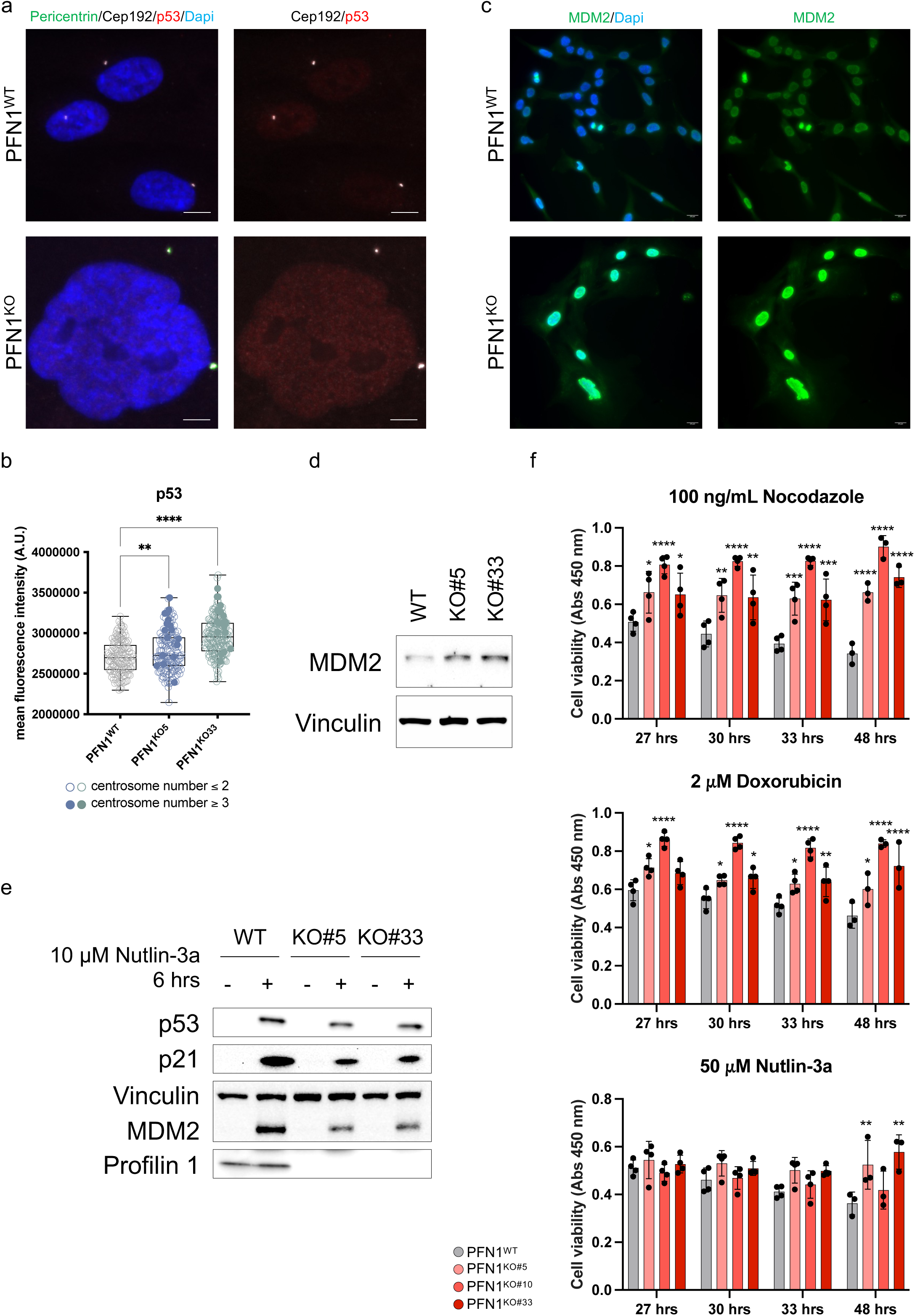
Profilin 1 loss affects the MDM2-p53-mediated stress response and confers resistance to genotoxic stress **a.** Representative immunofluorescence images of WT (top) and PFN1^KO^ (bottom) RPE1 cells stained for p53 (red) and centrosomal markers (Cep192, white; Pericentrin, green), with nuclei counterstained with DAPI (blue). Scale bar, 10 µm. **b.** Bar plot showing mean nuclear p53 fluorescence intensity in WT and PFN1^KO^ cells. Filled dots indicate cells containing three or more centrosomes (endoreplicated cells), which are distributed across the full range of p53 fluorescence intensities. Statistical analysis (ordinary one-way ANOVA) was performed on the total population of nuclei; \*\**p* = 0.0073; \*\*\*\**p* < 0.0001. *N* = 143 cells for WT; 161 cells for KO#5; 199 cells for KO#33. **c.** Representative immunofluorescence images of WT (top) and PFN1^KO^ (bottom) RPE1 cells stained for MDM2 (green) and counterstained with DAPI (blue). Scale bar, 20 µm. **d.** Western blot analysis of MDM2 expression in WT and two PFN1^KO^ RPE1 cells. Vinculin was used as a loading control. **e.** Western blot analysis of p53 expression in WT and PFN1^KO^ cells treated or not with Nutlin-3a (10 µM) for 6 hours. p21 and MDM2 were used as transcriptional readouts of p53 activation. Vinculin was used as a loading control. **f.** Cell viability assays (CCK-8) showing absorbance at 450 nm for WT and PFN1^KO^ cells following exposure to 100 ng/mL Nocodazole (top), 2 µM Doxorubicin (middle), and 50 µM Nutlin-3a (bottom), for the indicated durations. Statistical analyses were performed using 2way ANOVA; \*\*\*\**p* < 0.0001; \*\*\**p* < 0.001; \*\**p* < 0.005; \**p* < 0.05.

We next assessed whether this defective checkpoint response translated into altered drug sensitivity. Cell viability assays revealed that PFN1^KO^ cells were markedly more resistant to Doxorubicin and Nocodazole, and displayed late tolerance to Nutlin-3a (**Fig. 4f**). In contrast, Olaparib, a PARP inhibitor that targets cancer cells by exploiting deficiencies in homologous recombination (HR) repair^26^, did not show differential toxicity between WT and KO cells, suggesting that PFN1^KO^ cells do not exhibit a major defect in HR repair capacity (**Supplementary Fig. 4d**).

Finally, to investigate DNA damage signaling in basal conditions, we stained for γH2AX and 53BP1. PFN1^KO^ cells showed few γH2AX foci, consistent with low levels of acute damage, but a higher frequency of 53BP1 foci, indicative of persistent DNA lesions or past mitotic errors (**Supplementary Fig. 4e-g**; refs.^27,28^).

Together, these findings indicate that *PFN1* loss is associated with attenuated p53 activation and reduced responsiveness to chemotherapy-induced stress, creating conditions in which cells with abnormal genome content can persist and continue proliferating.

### 5. Attenuation of late cell cycle regulators upon *PFN1* loss

To gain insight into the molecular pathways affected by *PFN1* loss, we performed quantitative proteomic profiling of parental RPE1 cells and two independent PFN1 knock-out clones. Protein abundances were quantified across four biological replicates per condition, and differential analysis was carried out to identify proteins and pathways impacted by PFN1 depletion. Unsupervised principal component analysis revealed a clear separation between WT and Profilin 1–deficient samples (**Supplementary Fig. 5a**). Differential abundance analysis was performed independently for each KO clone relative to WT using a Welch’s t-test, applying thresholds of log₂ fold change ≥ 0.6 or ≤ −0.6 and *p* < 0.05. Using these criteria, we identified 1,175 downregulated and 78 upregulated proteins in one PFN1^KO^ clone, and 632 downregulated and 72 upregulated proteins in the second clone (**Fig. 5a**). While each clone exhibited both shared and clone-specific changes, a substantial subset of proteins and pathways were consistently deregulated in the same direction in both PFN1^KO^ clones, pointing to common PFN1-dependent programs (**Supplementary Data 2**; **Fig. 5b**). Strikingly, among the downregulated proteins, we observed a coordinated and pronounced attenuation of core regulators of mitotic entry and execution, including CDK1, PLK1 as well as additional mitotic effectors, indicating a selective shutdown of the mitotic program upon *PFN1* loss (**Fig. 5c**). This effect was not restricted to isolated proteins but included entire functional modules required for mitotic entry and progression (**Fig. 5c,d**). Indeed, proteins involved in spindle–kinetochore attachment and checkpoint control were significantly reduced, consistent with reduced mitotic competence and impaired fidelity of chromosome segregation. In parallel, we observed a broad depletion of the APC/C–E2 axis, including multiple APC/C subunits and the E2 enzymes UBE2C and UBE2S. Importantly, this proteomic signature cannot be explained by a mere enrichment of the bulk PFN1^KO^ population in G1. First, we detected a concomitant reduction of replication licensing factors (MCM2–7 and ORC2–4), whose expression peaks in G1 to enable origin relicensing^29^, arguing against a simple shift in population distribution toward G1 (**Fig. 5d**). In line with this, many G1- and S-phase proteins were not increased but remained unchanged in PFN1^KO^ cells (**Supplementary Fig. 5b,c**; **Methods**), further excluding a bulk enrichment in G1 and indicating comparable cell cycle distributions between conditions. Second, not all mitotic markers were uniformly depleted: we identified 49 well-characterized mitotic proteins that were comparably expressed between WT and PFN1^KO^ cells (**Fig. 5e**; **Supplementary Fig. 6a,b**; **Methods**), indicating that the observed changes reflect a specific mitotic bypass biochemical signature.

**Figure 5:**
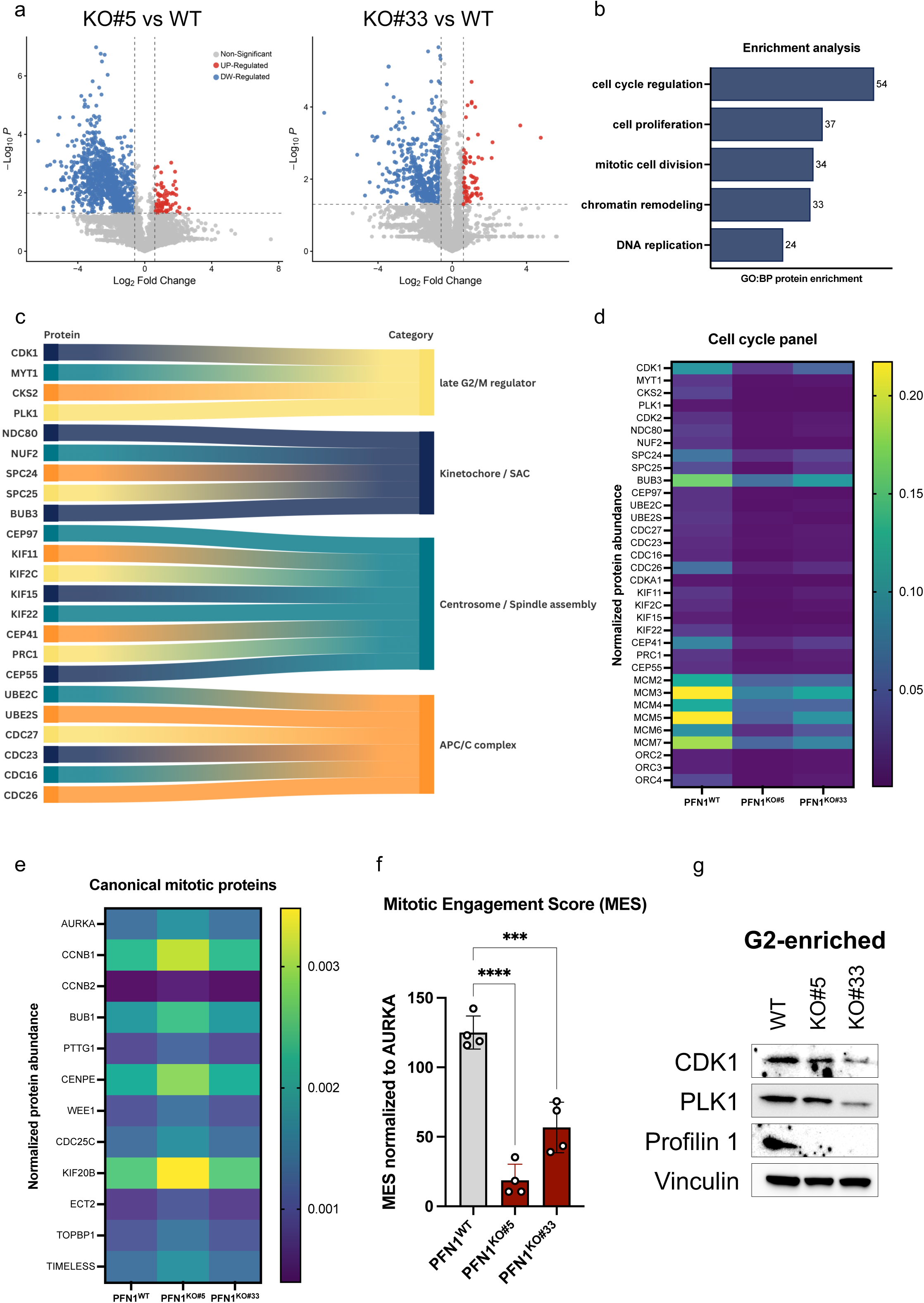
Coordinated deregulation of G2/M regulators upon PFN1 loss **a.** Volcano plots showing differential protein abundance between PFN1^KO#5^ and WT (left) and between PFN1^KO#33^ and WT RPE1 cells. Each dot represents one protein plotted according to log₂ fold change (x axis) and −log₁₀(*p*-value) (y axis). Proteins meeting the indicated significance thresholds are highlighted in blue (downregulated) or red (upregulated). **b.** Pathway enrichment analysis of proteins commonly downregulated in both PFN1^KO^ clones, performed using the g:Profiler tool. **c.** Alluvial plot showing the distribution of selected proteins across the functional categories of late G2/M regulators, kinetochore/spindle assembly checkpoint (SAC), centrosomes/spindle assembly, and APC/C complex. **d.** Heat map showing normalized protein abundance of selected cell cycle regulators in WT and PFN1^KO^ RPE1 cells. Protein expression levels are shown as row-scaled values across samples. **e.** Heat map showing normalized protein abundance of canonical mitotic proteins in WT and PFN1^KO^ RPE1 cells. Protein expression levels are shown as row-scaled values across samples. **f.** Bar graph showing the mitotic engagement score (MES) calculated in WT and PFN1^KO^ RPE1 cells as the ratio of the mean abundance of four selected mitotic regulators representing key stages of mitosis (CDK1, NDC80, PRC1, and UBE2C) to that of a stable mitotic protein (AURKA). Each dot represents an independent replicate. Statistical analysis was performed using ordinary one-way ANOVA; \*\*\*\**p* < 0.0001, \*\*\**p* = 0.0002. **g.** Western blot analysis of two G2/M regulators (CDK1 and PLK1) in G2-enriched WT and PFN1KO RPE1 cells, isolated as Cdt1^high^/Geminin^high^ FUCCI-expressing cells. Vinculin was used as a loading control.

To quantitatively assess the extent of mitotic commitment, we computed a ***mitotic engagement score*** (**MES**) based on the combined abundance of selected mitotic regulators representing key stages of the mitotic process. Specifically, the score integrates: CDK1 (late G2/M transition), NDC80 (kinetochore function), PRC1 (spindle organization), and UBE2C (APC/C-dependent mitotic progression), normalized to AURKA as a relatively stable mitotic reference (**Fig. 5e**). The MES revealed a progressive reduction in mitotic engagement across PFN1^KO^ clones compared to WT cells (**Fig. 5f**), indicating an impairment to fully activate the mitotic program.

To functionally validate these proteomic findings in a cell cycle–resolved manner, we isolated G2-enriched populations from FUCCI-expressing WT and PFN1^KO^ RPE1 cells, by FACS-sorting Cdt1^high^/Geminin^high^ cells. Immunoblot analysis of these purified populations confirmed a marked reduction in the key mitotic regulators CDK1 and PLK1 in PFN1^KO^ cells compared to WT (**Fig. 5g**).

Together, these findings support a model in which *PFN1* loss impairs the activation of the mitotic program prior to mitotic entry, thereby predisposing cells to bypass mitosis.

### 6. Profilin 1 depletion increases the metastatic potential of osteosarcoma cells

To assess the contribution of *PFN1* loss to tumor progression in vivo, we performed xenograft experiments in immunodeficient mice. Because RPE1 cells are non-tumorigenic irrespective of *PFN1* status, they were not suitable for in vivo studies. We therefore transitioned to a cancer-relevant model and selected osteosarcoma cells, a tumor type in which *PFN1* is frequently downregulated and/or deleted (**refs.**^11,15^**; Supplementary Fig. 7a,b**) and that exhibits one of the highest rates of whole-genome doubling^10^. This approach allowed us to test the hypothesis that *PFN1* loss does not act as a transforming event *per se*, but rather licenses genome doubling, chromosomal instability, and pro-tumorigenic adaptations within an oncogenic context. SaOS2 cells were selected due to their aggressive and metastatic behavior and because they display relatively low basal levels of Profilin 1 (**Supplementary Fig. 7a, right**). To further reduce Profilin 1 expression, we generated two independent homozygous *PFN1* knockout clones (#4 and #5) using CRISPR/Cas9-mediated genome editing (**Supplementary Fig. 7c,d**). For in vivo experiments, PFN1-deficient cells were injected as a 1:1 mixture of the two KO clones to minimize potential clone-specific effects. Parental SaOS2 cells or PFN1^KO^ cells were injected intratibially into NOD-SCID-Gamma (NSG) mice, and tumor growth was monitored for 6 weeks, after which animals were sacrificed for macroscopic and organ-level analysis (**Fig. 6a**). Given the limited cohort size (n = 2 mice per condition), these experiments were designed as a proof-of-concept to explore the potential impact of *PFN1* loss on osteosarcoma progression and dissemination. At the time of sacrifice, primary tumors arising from parental and PFN1^KO^ SaOS2 cells were comparable in size (**Fig. 6b,c**), indicating that *PFN1* loss does not markedly enhance local tumor growth in this setting. In contrast, mice injected with PFN1-deficient cells exhibited a higher metastatic burden compared to controls. While parental SaOS2 tumors showed either no or limited metastatic dissemination, PFN1^KO^ tumors were consistently associated with the presence of multiple metastatic nodules, particularly in the liver and lungs (**Fig. 6c,d**). Notably, PFN1^KO^-injected mice exhibited an approximately twofold increase in the frequency of lung metastases compared to parental controls (**Fig. 6e**). Histological analysis of lung sections further revealed that metastatic PFN1^KO^ SaOS2 cells frequently displayed enlarged nuclei (**Fig. 6e, bottom**), consistent with the presence of polyploid cells within metastatic lesions.

**Figure 6:**
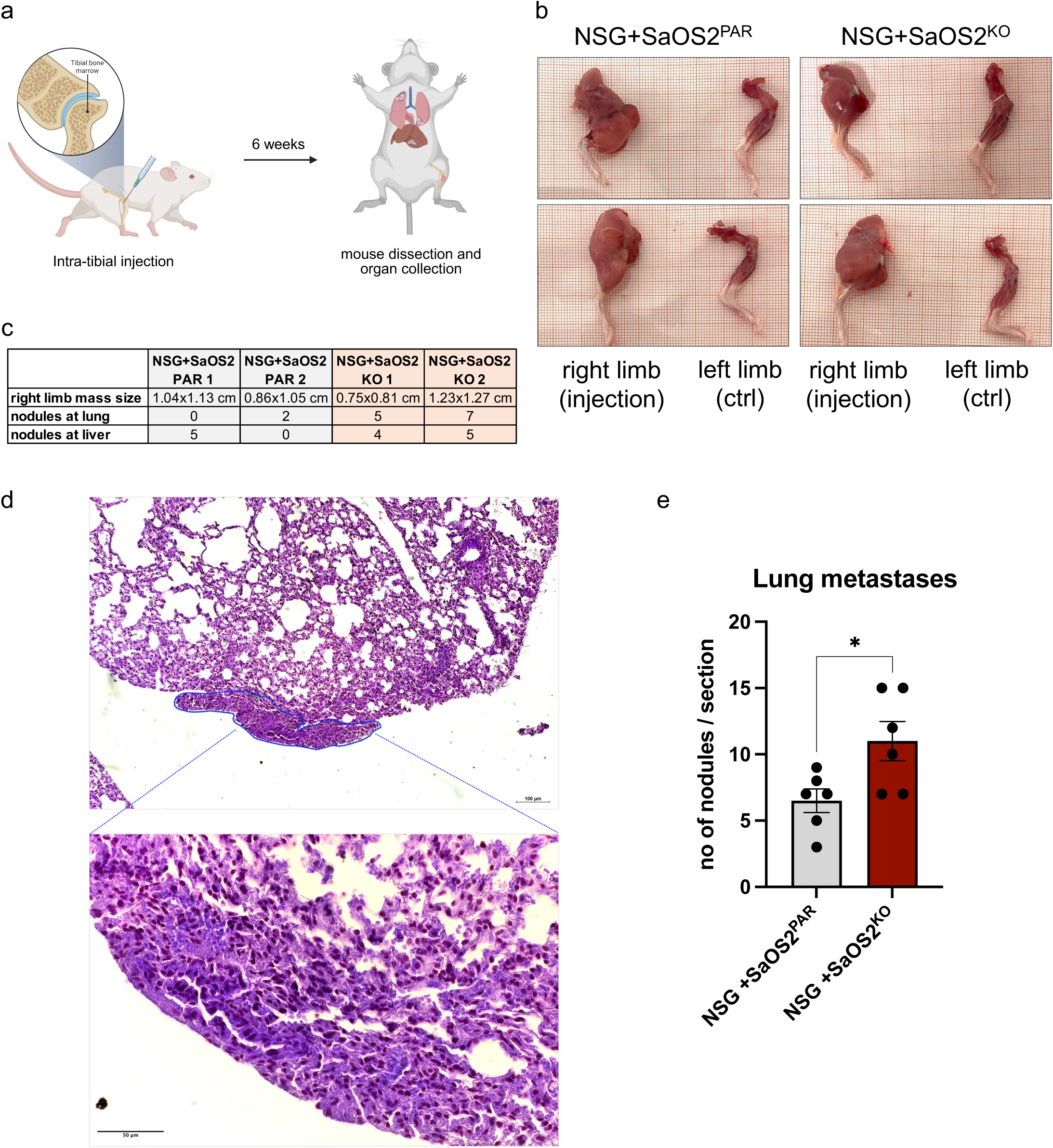
Profilin 1 depletion increases the metastatic potential of osteosarcoma cells **a.** Schematic overview of the intratibial injection of SaOS2 cells into the bone marrow cavity of NOD-SCID-Gamma (NSG) mice and subsequent tissue collection at endpoint. **b.** Representative photographs of injected and contralateral non-injected hind limbs of NSG mice showing osteosarcoma growth at the joint region. **c.** Table summarizing osteosarcoma mass size and the number of metastatic nodules detected in the lungs and liver of NSG mice injected with parental (PAR) or PFN1^KO^ SaOS2 cells. **d.** Representative hematoxylin and eosin (H&E)–stained lung section showing normal alveolar architecture and infiltration by metastatic osteosarcoma cells (blue contour). Bottom, higher-magnification view of the metastatic lesion. **e.** Quantification of the number of metastatic sites detected across lung sections from NSG mice injected with parental or PFN1 KO SaOS2 cells.

Although exploratory, these in vivo findings suggest that PFN1 depletion promotes tumor dissemination. This observation is consistent with previous evidence indicating that genome-doubled and polyploid states can enhance cellular plasticity and adaptability, features that have been associated with tumor progression and metastatic potential^9,30^.

## Discussion

Whole-genome doubling (WGD) is a frequent event in cancer evolution and is associated with genomic instability, tumor progression, and poor clinical outcome^2–4^. However, the cellular mechanisms that enable genome doubling in p53-proficient cells remain incompletely understood. In this study, we identify loss of Profilin 1 (PFN1) as a driver of genome doubling through a mechanism involving mitotic bypass and endoreplication, and we provide evidence that this process contributes to progressive polyploidization, chromosomal instability, and enhanced cellular plasticity.

Our data indicate that PFN1 deficiency destabilizes the commitment to mitotic entry. Live-cell imaging revealed that a substantial fraction of PFN1-deficient cells fails to initiate mitosis, instead transitioning directly into a G1-like state without chromosome segregation. These cells subsequently re-enter S phase and replicate their DNA, giving rise to mononucleated tetraploid/polyploid cells. This behavior is consistent with mitotic bypass followed by endoreplication and supports the notion that *PFN1* loss disrupts the regulatory mechanisms governing the G2/M transition. In this context, PFN1 appears to function as a gatekeeper of mitotic commitment. Our EdU incorporation analyses further suggest that mitotic bypass in PFN1-deficient cells is frequently followed by incomplete or inefficient DNA replication, as indicated by reduced EdU signal intensity compared with canonical S phase. This observation is consistent with emerging evidence that Profilin 1 directly participates in replication fork progression and origin firing, and that its loss leads to replication stress and under-replication of genomic regions^14^. In this context, the attenuated EdU incorporation observed in our system likely reflects compromised replication dynamics rather than an absence of replication *per se*. Consistent with this interpretation, PFN1-deficient cells displayed increased 53BP1 foci despite relatively low levels of acute DNA damage. Persistent 53BP1 positivity is commonly associated with chronic or unresolved DNA lesions, including under-replicated regions transmitted across cell cycles^28^. Together, these findings support a model in which *PFN1* loss induces a mitotic bypass that does not behave as a terminal fail-safe response. In principle, exit from G2 without mitosis could represent a protective mechanism if followed by stable cell cycle arrest^31–33^. However, our data show that PFN1-deficient cells can re-enter S phase after bypass and, in some cases, subsequently attempt aberrant mitotic divisions. Thus, *PFN1* loss converts mitotic bypass from a potential barrier to proliferation into a permissive route toward WGD and continued propagation of genomically abnormal cells. Mechanistically, our proteomic analyses revealed a coordinated downregulation of late cell cycle and mitotic regulators, including CDK1 and PLK1. These findings support the existence of a partially suppressed mitotic program that may prevent full activation of mitotic entry while allowing continued cell cycle progression. Such an altered regulatory landscape is consistent with the prolonged G2 phase and defective checkpoint execution observed in PFN1-deficient cells. One possible interpretation is that *PFN1* loss does not simply delay cells before an otherwise normal mitosis, but lowers the biochemical threshold required to maintain full mitotic competence. In this scenario, reduced abundance of late G2/M and mitotic regulators would favor an unstable pre-mitotic state in which cells can initiate G2 but fail to irreversibly commit to mitotic entry. Reduced CDK1 abundance in PFN1^KO^ endoreplicating cells is consistent with a conserved mechanism promoting mitotic bypass and genome doubling. In multiple developmental contexts, downregulation of CDK1 activity is required for the transition from mitotic cycles to endoreplication^34–36^. Moreover, pharmacological inhibition of CDK1 has been shown to induce endoreplication and polyploidization in mammalian systems^20^. Together, these findings suggest that the decrease in CDK1 observed in PFN1-deficient cells may reflect acquisition of a permissive endoreplicative state. This model is further supported by the marked resistance of PFN1^KO^ cells to nocodazole treatment. Nocodazole disrupts spindle assembly and primarily affects cells that enter mitosis by inducing prolonged mitotic arrest, which can lead to cohesion fatigue, mitotic slippage, or cell death^37,38^; however, PFN1-deficient cells display high tolerance to this treatment, consistent with their reduced reliance on canonical mitotic entry. Mitotic bypass thus represents not only a route to genome doubling but also a mechanism of escape from therapies targeting mitotic processes.

Another key observation emerging from this work is that *PFN1* loss enables genome doubling in the presence of functional p53. This finding addresses a long-standing paradox in cancer biology, as WGD is frequently detected in tumors retaining wild-type TP53. In PFN1-deficient cells, we observed increased nuclear accumulation of MDM2 and a blunted p53 response following genotoxic stress, suggesting functional attenuation of the p53 surveillance pathway. Importantly, single-cell whole-genome sequencing excluded copy-number gains at the MDM2 locus, ruling out clonal selection or genomic amplification as the cause of MDM2 upregulation. This aspect is particularly important because the proliferative outcome of mitotic bypass is expected to depend on the ability of checkpoint pathways to recognize and restrain cells with abnormal genome content^39^. In PFN1-deficient cells, p53 is detectable but fails to scale appropriately with nuclear size, centrosome amplification, or replication-associated stress. Although the precise molecular mechanisms underlying this deregulation remain to be defined, these results indicate that PFN1 deficiency can uncouple genome doubling from canonical p53-mediated cell cycle arrest and apoptosis. Such functional attenuation of checkpoint signaling may allow cells to tolerate replication-associated damage and proceed through additional rounds of DNA synthesis. In this regard, it is noteworthy that recent work has shown that p53 activity can, in certain contexts, be required for mitotic bypass to occur, rather than strictly acting as a barrier to it^31^. Dysregulated MDM2 accumulation may therefore skew p53 responses toward permissive states that allow mitotic bypass and genome reduplication.

Notably, a conserved link between Profilin 1 and endoreplication has been described in *Drosophila*, where reduced activity of the profilin homolog chickadee was shown to promote premature entry into endoreplication of salivary gland cells, leading to aberrant polyploidization^40^. Although observed in a developmental setting, these findings parallel our results in human cells and support the idea that profilins function constitutes an evolutionarily conserved constraint on genome reduplication.

The in vivo experiments presented here provide preliminary support for the biological relevance of this mechanism. Mice injected with PFN1-deficient osteosarcoma cells displayed a higher frequency of pulmonary metastatic lesions compared with controls, and disseminated tumor cells frequently exhibited enlarged nuclei compatible with a polyploid state. Although these observations remain exploratory, they are consistent with the idea that genome-doubled and polyploid cells possess increased phenotypic plasticity, which may facilitate survival during dissemination and colonization of distant tissues ^9,30^. We speculate that the cellular plasticity enabled by *PFN1* loss (i.e., mitotic bypass, replication stress tolerance, and checkpoint evasion) may provide a selective advantage during the complex and hostile process of dissemination and colonization of distant organs.

Altogether, our observations support a model in which *PFN1* loss affects two complementary surveillance mechanisms of genome integrity: (i) first, attenuation of late G2/M and mitotic regulators weakens mitotic commitment, increasing the probability that cells exit G2 without chromosome segregation; (ii) second, defective scaling of the p53 response prevents these genome-doubled cells from undergoing cell cycle arrest. The combination of impaired mitotic commitment and checkpoint evasion therefore creates a permissive state in which mitotic bypass fuels WGD and chromosomal instability. In conclusion, our study positions Profilin 1 as a critical regulator of cell cycle fidelity whose loss destabilizes late cell cycle control, enabling mitotic bypass, endoreplication, and whole-genome doubling in p53-proficient cells. These findings provide new insight into the cellular logic of genome doubling and suggest that PFN1-dependent pathways may represent vulnerabilities in polyploid and genomically unstable cancers.

## Materials and Methods

### Copy-number and whole-genome doubling analysis

Whole-genome doubling (WGD) status for TCGA Pan-Cancer Atlas samples was determined using genome doublings values reported in the ABSOLUTE-derived file TCGA_mastercalls.abs_tables_JSedit.fixed.txt (GDC PanCanAtlas; https://gdc.cancer.gov/about-data/publications/pancanatlas). Samples with genome doublings ≥1 were classified as WGD-positive, those with genome doublings equal to zero as WGD-negative, and cases lacking WGD information were excluded. Gene-level copy-number variation (CNV) data for PFN1 were obtained from the TCGA PanCanAtlas GISTIC2 thresholded dataset (Broad Institute/UCSC Xena; DOI: 10.5281/zenodo.827323). GISTIC2 values of −2 and −1 were classified as PFN1 deletion (PFN1del), corresponding to deep/homozygous and shallow/heterozygous deletion, respectively, whereas 0 was classified as PFN1 copy-number wild-type/neutral (PFN1wt). TCGA barcodes were harmonized and intersected across datasets to retain matched samples with both WGD and PFN1 CNV information. Statistical analysis of the association between *PFN1* deletion and WGD status was performed using two-sided Fisher’s exact tests on 2×2 contingency tables for each tumor type. Odds ratios (OR) and 95% confidence intervals (CI) were calculated using the log-Woolf method with continuity correction when necessary. P-values were adjusted for multiple testing using the Benjamini–Hochberg false discovery rate (FDR) method. For pan-cancer analysis, contingency tables were pooled across tumor types. To assess whether the association between WGD and *PFN1* deletion was independent of tumor context, multivariable logistic regression models were fitted at the sample level with WGD status as predictor and tumor type included as a covariate. All statistical analyses were performed using R.

### Cell culture and treatments

hTERT-RPE1 (RPE1) were cultured in DMEM/F12 medium supplemented with 10% fetal bovine serum (FBS) and 1% penicillin–streptomycin (P/S), and maintained at 37 °C in a humidified atmosphere with 5% CO2. Parental RPE1 cells were used as wild-type (WT) controls. *PFN1* knockout (PFN1^KO^) RPE1 cell lines were generated previously at passage 7 [ref. ^15^] and two independent knockout clones (KO#5 and KO#33) were used throughout the study, unless otherwise indicated, as in **Fig. 4f** where KO#10 was also included. To assess the temporal dynamics of polyploidization, RPE1 WT and PFN1^KO^ cells were cultured and analyzed at different passage ranges: early passages (p<15), mid passages (p=20–30), and late passages (p>40). Cells were routinely cryopreserved at each passage interval to ensure reproducibility across experiments. RPE1 cells stably expressing the FUCCI reporters were cultured under the same conditions and analyzed at mid passages.

Human osteosarcoma SaOS2 cells were cultured in RPMI 1640 medium supplemented with 10% FBS and 1% P/S, and maintained at 37 °C with 5% CO₂.

All cell lines were tested for mycoplasma contamination (PCR Mycoplasma Detection Kit, Applied Biological Materials, G238) and were found to be negative. Cell lines were not authenticated.

For pharmacological induction of p53 signaling, cells were treated with Nutlin-3a (Sigma-Aldrich, SML0580; 10 µM) for 6 h, or with Doxorubicin (Sigma-Aldrich, D1515; 2 µM) for 2 h, or with hydroxyurea (HU) (SelleckChem, 50-136-1749; 2 mM) overnight (17 h). For cytotoxicity assays, cells were treated with Doxorubicin (Sigma-Aldrich, D1515; 2 µM), Nutlin-3a (Sigma-Aldrich, SML0580; 50 µM), Nocodazole (Sigma-Aldrich, M1404; 100 ng/mL), or Olaparib (SelleckChem, S1060; 40 µM) for the durations indicated in the corresponding figure legends.

### Cell viability assay (CCK-8)

Cell viability was assessed using the Cell Counting Kit-8 (CCK-8; Dojindo, CK04) assay according to the manufacturer’s instructions. Briefly, 2,000 cells per well were seeded in 96-well plates and allowed to adhere overnight. Cells were then treated as described above. At the specified time points, CCK-8 reagent was added to each well and plates were incubated at 37 °C for the time indicated in the figures. Absorbance was measured at 450 nm using a microplate reader. To account for potential differences in initial cell seeding, absorbance values were normalized to the 3-h time point of the CCK-8 assay for each condition.

### Generation of RPE1-FUCCI stable cell lines

The tFUCCI(CA)2-pCSII-EF lentiviral vector^20^ was used to generate RPE1 cell lines stably expressing the FUCCI cell cycle reporters. The plasmid was first transformed into NEB® 5-alpha Competent E. coli (DH5α, high efficiency), and a single bacterial colony was selected, expanded, and used for plasmid amplification and purification according to standard protocols. Lentiviral particles were produced in HEK293T cells by transient transfection. Briefly, HEK293T cells were transfected with 4 µg of tFUCCI(CA)2-pCSII-EF plasmid, 4 µg of pMD2.G (Addgene #12259), and 4 µg of psPAX2 (Addgene #12260) using FuGENE HD Transfection Reagent (Promega, E2311) in OptiMEM medium (Thermo Fisher Scientific, 51985034). Cells were incubated at 37 °C in a humidified atmosphere with 5% CO₂ for 16 h, after which the transfection medium was replaced with 6 mL of fresh OptiMEM. Viral supernatants were collected 24 h later and filtered through a 0.45 µm filter to remove cellular debris. RPE1 WT and PFN1^KO^ cells (p=20) were incubated with filtered viral particles at 37 °C with 5% CO₂ for 24 h. Following infection, GFP- and RFP-positive cells were isolated by fluorescence-activated cell sorting (FACS) using a Sony SH800 cell sorter (BD FACSDiva software, version 8.0.1).

FUCCI-expressing RPE1 cell lines were established as bulk populations rather than single-cell–derived clones to minimize clonal selection bias and preserve population-level heterogeneity.

### Generation of RPE1-H2B-mCherry stable cell lines

To visualize chromatin dynamics during live-cell imaging, RPE1 cells were engineered to stably express histone H2B fused to mCherry. Lentiviral particles were produced in HEK293T cells by transient transfection with 4 µg pLenti6-H2B-mCherry plasmid (Addgene #89766), 4 µg pMD2.G (Addgene #12259), and 4 µg psPAX2 (Addgene #12260), using the same transfection conditions described above for FUCCI virus production. Following infection, mCherry-positive RPE1 PFN1^WT^ and PFN1^KO^ cells were isolated by FACS using a Sony SH800 cell sorter (BD FACSDiva software, version 8.0.1). Cell lines expressing H2B–mCherry were established as bulk populations.

### SaOS2 CRISPR/Cas9-mediated *PFN1* knockout

*PFN1* knockout SaOS2 cell lines were generated using the same CRISPR/Cas9 strategy previously employed for RPE1 *PFN1* knockout cells^15^. Briefly, the CRISPR guide RNA (gRNA) sequence 5′-CACCGTTCGTACCAAGAGCACCGGT-3′, targeting the *PFN1* locus, was designed and annealed oligonucleotides were cloned into the pSpCas9n(BB)-2A-GFP plasmid (Addgene, PX458). SaOS2 cells were transfected with the CRISPR/Cas9 construct using Lipofectamine 2000 (Invitrogen), according to the manufacturer’s instructions. Forty-eight hours after transfection, GFP-positive cells were isolated by fluorescence-activated cell sorting (FACS) and seeded into 96-well plates to obtain single-cell–derived colonies. Candidate *PFN1* knockout clones were screened by Sanger sequencing to confirm the presence of frameshift-inducing mutations at the target locus and by western blotting to verify loss of Profilin 1 protein expression. Potential off-target mutations in genes predicted by the Optimized CRISPR Design Tool (http://crispr.mit.edu/), including GRAMD4 and INTS6, were excluded by Sanger sequencing. Validated homozygous *PFN1* knockout clones were expanded and used for subsequent in vitro and in vivo experiments. Where indicated, independent knockout clones were combined at a 1:1 ratio to minimize potential clone-specific effects.

### Cell cycle analysis by propidium iodide staining and serum starvation

To accurately assess cellular ploidy and avoid confounding contributions from cells in G2/M phase, RPE1 WT and PFN1^KO^ cells were synchronized by serum starvation prior to DNA content analysis. This approach allows discrimination between *bona fide* polyploid cells and diploid cells transiently in G2/M. Cells were cultured in complete medium and subsequently serum-starved for a total of 72 h: after the first 24 h of starvation, the medium was replaced with fresh serum-free medium and cells were maintained under serum-free conditions for the remaining 48 h prior to fixation. Following serum starvation, cells were harvested, washed with cold phosphate-buffered saline (PBS), and fixed in ice-cold 100% methanol. Fixed cells were incubated overnight at −20 °C. For flow-cytometry analysis, cells were resuspended in PBS containing RNase A (0.2 mg/mL) and incubated for 1 h at 37 °C to remove RNA contamination. Cells were then stained with propidium iodide (PI, 50 µg/mL) for 30 min at room temperature, protected from light. DNA content was analyzed by flow cytometry using a BD FACSCanto II instrument (Becton Dickinson). For cell cycle profiles and ploidy distributions, 10’000 events were recorded for each FACS analysis, and the data were analysed using FlowJo software (v10.10.0). The 2N DNA content peak was defined based on serum-starved WT RPE1 cells and used as reference for ploidy quantification across all samples. Ploidy measurements were performed at early (p<15) and late (p>40) passages in three independent biological experiments.

### Single-cell whole-genome sequencing

Single-cell whole-genome sequencing (scWGS) was performed on early-passage (p<15) WT and PFN1^KO^ RPE1 cells to assess DNA copy-number states at single-cell resolution. To avoid confounding effects from cells in G2/M phase, cells were serum-starved as described above prior to analysis. Following serum starvation, cells were harvested, pelleted, and processed for single-cell genome sequencing. In parallel, an aliquot of the same cell population was subjected to EdU incorporation assays (described below) to verify effective cell cycle arrest in G0/G1 and to exclude the presence of cells undergoing S phase. This control ensured that cells displaying 4N DNA content reflected *bona fide* tetraploid G1 cells rather than diploid cells in G2 phase.

Single-cell libraries were sequenced on a NextSeq 2000 platform (Illumina; up to 77 cycles). Aligned sequencing reads (BAM files) were analyzed using the developer version of AneuFinder (version 1.7.4; from GitHub). The ground ploidy for these samples was constrained between 3.5 and 4.5 (parameters: min.ground.ploidy and max.ground.ploidy; samples with whole-genome duplication). Following GC correction and blacklisting of artefact-prone regions (extreme low or high coverage in control samples), libraries were analyzed using the dnacopy and edivisive copy number calling algorithms with variable width bins (average binsize = 1 Mb; step size = 500 kb) and breakpoint refinement (confint = 0.95). Both the blacklists and variable width bins were constructed using a euploid reference^41^. Results were curated by requiring a minimum concordance of 90% between the results of the two algorithms. Libraries with an average of fewer than 10 reads per chromosome copy per bin (2-somy: 20 reads, 3-somy: 30 reads, etc.) were discarded.

For each bin, the aneuploidy score was calculated as the absolute difference between the observed and euploid copy number. The weighted average of the bins is the score per library, which was averaged to the score per sample. The heterogeneity score per bin was calculated as the proportion of pairwise comparisons (cell 1 vs. cell 2, cell 1 vs cell 3, etc.) that showed a difference in copy number (e.g. cell 1: 2-somy, cell 2: 3-somy). The score of each sample is the weighted average of the scores of all bins.

The data were demultiplexed using sample-specific barcodes and changed into fastq files using bcl2fastq (Illumina; version 1.8.4). Reads were aligned to the human reference genome (GRCh38/hg38) using Bowtie2 (version 2.2.4; ref. ^42^) and duplicate reads were marked with BamUtil (version 1.0.3; ref. ^43^).

### EdU incorporation assay and quantification

DNA synthesis was assessed by EdU incorporation using the Click-iT™ EdU Imaging Kit (Thermo Fisher Scientific, C10338), according to the manufacturer’s instructions. EdU was added to the culture medium at a final concentration of 10 µM for either 30 min (**Supplementary Fig. 2b**) or 18 h (**Fig. 3e** and **Supplementary Fig. 3d**). Following EdU labeling, cells were fixed and subjected to the Click-iT reaction for 15 min to fluorescently label incorporated EdU. Nuclei were counterstained as indicated. Images were acquired using an upright widefield microscope (Leica DM6B, Leica Microsystems) equipped with a motorized XY stage and a 40× objective, using MetaMorph software (version 7.10.1, Molecular Devices). For each field, z-stacks of fluorescence images were automatically collected at 0.5 µm intervals. Images were processed using ImageJ software and displayed as maximum intensity projections. EdU fluorescence intensity was quantified on maximum intensity projections by manually outlining individual nuclei using the wand selection tool in ImageJ.

For Edu Incorporation in **Fig. 3f** and **Supplementary Fig. 3e**, cells were incubated with EdU (10 μM, Thermo Fisher Scientific, C10640) for 4 hours, fixed with cold methanol, washed, and stained with Alexa-Fluor Azide 647 click reagent. EdU stained cells were then washed in 1x PBS and stained with PI (50 μg/mL) in the presence of RNase A (0.1 mg/mL) and incubated for 30 minutes (37 °C, dark), and then washed. The DNA Content was measured on a BD FACSCanto II instrument (Becton Dickinson).

### Image analysis and quantification

Image analysis and quantitative measurements were performed using ImageJ software (NIH). Nuclear area, DAPI, p53, MDM2, and EdU fluorescence intensities were quantified on maximum intensity projections by manually outlining individual nuclei using the wand selection tool. Binucleated and multinucleated cells were identified and manually counted based on nuclear morphology. Centrosomes were defined and quantified exclusively as structures positive for both Cep192 and Pericentrin immunostaining. A threshold of 250 µm² was used to define normal-sized nuclei, corresponding to the mean nuclear area measured in asynchronous *PFN1* wild-type RPE1 cultures.

For live-cell imaging analyses, custom Fiji macros were used to extract quantitative information from FUCCI- and H2B-mCherry–expressing cells. FUCCI-based cell cycle tracking was performed using a dedicated macro to assign cell cycle phases and transitions over time. Mitotic progression and mitotic outcomes were analyzed on H2B-mCherry time-lapse movies using a customized mitosis analysis pipeline, enabling annotation of mitotic entry, mitotic exit, abnormal divisions, chromosome segregation defects, and cell fate.

For phase duration measurements in FUCCI time-lapse movies, only cells for which both the onset and completion of the respective cell cycle phase were captured within the imaging window were included in the analysis. Cells that entered or exited a given phase outside the recording interval were excluded.

DNA damage quantification was performed using a custom Fiji macro that automatically detected and quantified nuclear DNA damage signals, including γH2AX and 53BP1 foci, based on fluorescence intensity and spot coverage within nuclear regions. Fluorescence images were analyzed on z-projection stacks following intensity thresholding. For each nucleus, nuclear size and DAPI intensity were measured, together with the number of γH2AX or 53BP1 foci. Quantitative analyses were performed using identical thresholding and analysis parameters across all experimental conditions.

For long-term live-cell imaging, cells were manually centered using the Fiji Manual Tracking plugin to compensate for stage drift and cell movement. Image stacks were subsequently realigned using a custom macro to generate translated stacks prior to quantitative analysis. All image processing and quantifications were performed on raw data using identical analysis parameters across experimental conditions.

For histological analysis of the metastatic burden in the lung, the areas that contained nodules foci or micro-metastasis were classified manually and counted based on alterations of the lung tissue, which included *i*) clusters of morphologically altered epithelial cells in perivascular regions, and *ii*) localized remodeling of the alveolar regions, characterized by expansion or effacement of the alveolar septa by aggregates of tumor cells. The number of metastatic foci was calculated as the mean of nodules identified across three sections, ensuring a minimum of 50 µm between analyzed sections. Overall, the analyzed sections spanned approximately 200 µm of tissue depth, to minimize the risk of counting the same lesion multiple times, although not all intervening sections were examined.

### Time-lapse microscopy

For live-cell imaging experiments, 20,000 cells were plated on glass-bottom dishes (Dutscher, 627870). Time-lapse imaging was performed using a spinning disk microscope equipped with a motorized XY stage mounted on an inverted Nikon microscope. Images were acquired using a 20× dry objective and an sCMOS camera. All live-cell imaging experiments were performed in a temperature- and CO₂-controlled environment at 37 °C and 5% CO₂. Laser power and acquisition settings were optimized to achieve the best signal-to-background ratio while minimizing phototoxicity. A mild degree of phototoxic stress was nonetheless observed during the final hours of prolonged FUCCI time-lapse experiments, as indicated by occasional stalling of cells in G1 phase, and was taken into account during data interpretation. Multi-dimensional acquisitions were performed using MetaMorph software. Z-stacks of fluorescence images were automatically collected at 0.5 µm intervals. To ensure image stability, the initial frames acquired immediately after microscope equilibration were excluded from analysis, as minor focus drift associated with thermal stabilization was observed. Images were processed using ImageJ software and are shown as maximum intensity projections generated from deconvolved image stacks using an extension of MetaMorph. FUCCI time-lapse experiments were performed acquiring one image every 15 minutes for a total duration of 65–72 h, whereas H2B–mCherry live-cell imaging was carried out for 14–18 h. Time-lapse movies were exported using ImageJ software at a frame rate of 7 frames per second (fps).

### Indirect immunofluorescence

Cells were seeded on glass coverslips in 24-well plates and processed for immunofluorescence staining. Cells were fixed and permeabilized using 4% paraformaldehyde (Electron Microscopy Sciences, 15710) supplemented with 0.1% Triton X-100 in PBS for 20 min at 4 °C. Following fixation, cells were washed three times with PBS-T (PBS containing 0.1% Triton X-100 and 0.02% sodium azide) and blocked in PBS-T supplemented with 1% bovine serum albumin (BSA) for 30 min at room temperature. After blocking, coverslips were incubated with primary antibodies diluted in PBS-T containing 1% BSA for 1 h at room temperature. Cells were then washed three times with PBS-T/BSA and incubated with fluorophore-conjugated secondary antibodies diluted in PBS-T/BSA for 45 min at room temperature. Following secondary antibody incubation, cells were washed twice with PBS and counterstained with DAPI (3 µg/mL; Sigma-Aldrich, D8417) for 15 min at room temperature. Coverslips were subsequently washed twice with PBS and mounted using an antifade mounting medium.

Primary antibodies were used at the following dilutions: guinea pig anti-CEP192 (1:1200; R.B. laboratory), rabbit anti–β-catenin (1:250; Sigma-Aldrich, C2206), mouse anti–phospho-γH2AX (Ser139) (1:800; Abcam, ab22551), mouse anti-53BP1 (1:250; Millipore, MAB3802), rabbit anti-p53 (1:250; Abcam, ab17990), mouse anti-Pericentrin (1:500; Abcam, ab28144), mouse anti-PCNA (1:250; Santa Cruz Biotechnology, sc-56), and mouse anti-MDM2 (1:250; Invitrogen, MA1-113).

Secondary antibodies were used at a dilution of 1:250 and included: goat anti–guinea pig IgG (H+L) Alexa Fluor 647 (Thermo Fisher Scientific, A21450), goat anti–mouse IgG (H+L) Alexa Fluor 546 (Thermo Fisher Scientific, A11003), goat anti–rabbit IgG (H+L) Alexa Fluor 546 (Thermo Fisher Scientific, A11035), goat anti–mouse IgG (H+L) Alexa Fluor 488 (Thermo Fisher Scientific, A11001), and goat anti–rabbit IgG (H+L) Alexa Fluor 488 (Thermo Fisher Scientific, A11008).

### Protein extraction and western blotting

Total cell lysates were prepared using RIPA buffer supplemented with protease inhibitors (1:100; Applied Biological Materials, G135) and phosphatase inhibitors (1:100; Sigma-Aldrich, P0044). For osteosarcoma tissue samples, mechanical disruption and homogenization were performed using a TissueLyser LT (Qiagen) for 5 min at 50 Hz. Protein concentrations were determined using a BCA assay according to the manufacturer’s instructions. Protein lysates were denatured by heating at 85 °C for 3 min and separated by SDS–PAGE using 8–16% Tris–Glycine gels (Invitrogen) at 180 V for 30 min. Proteins were transferred onto nitrocellulose membranes using an iBlot transfer system (Invitrogen) for 5 min. Membranes were blocked for 1 h at room temperature in TBST (1× TBS, 0.05% Tween-20) containing 4% (w/v) non-fat dry milk. Membranes were incubated with primary antibodies diluted in blocking buffer, including: rabbit anti–Profilin 1 (Invitrogen, PA5-17444; 1:5000), mouse anti-p53 (Cell Signaling Technology, 2524; 1:1000), mouse anti-CHK1 (Santa Cruz Biotechnology, sc-8408; 1:200), rabbit anti–phospho-CHK1 (Ser317) (Cell Signaling Technology, 2344S; 1:500), rabbit anti-CHK2 (Cell Signaling Technology, 2662; 1:1000), rabbit anti–phospho-CHK2 (Thr68) (Cell Signaling Technology, 2661; 1:500), rabbit anti–phospho-Histone H2A.X (Ser139) (Cell Signaling Technology, 2577; 1:1000), mouse anti–PLK1 (Abcam ab-17056; 1:1000), mouse anti–CDK1 (BD Biosciences, BD-610038; 1:1000), mouse anti–MDM2 (Thermo Fisher Scientific, MA1-113; 1:500), rabbit anti–p21 (Cell Signaling Technology, 2947; 1:1000), rabbit anti–GAPDH (Sigma-Aldrich, G9545; 1:5000), mouse anti–α-Tubulin (Sigma-Aldrich, T6074; 1:15000), rabbit anti–Vinculin (Cell Signaling Technology, 13901; 1:10’000).

Following primary antibody incubation, membranes were washed and incubated with HRP-conjugated secondary antibodies for 1 h at room temperature. Protein signals were detected using enhanced chemiluminescence (SuperSignal™ West Pico Chemiluminescent Substrate, Thermo Scientific, 34080; or SuperSignal™ West Femto Maximum Sensitivity Substrate, Thermo Scientific, 34096) and visualized using a ChemiDoc imaging system (Bio-Rad). Equal protein loading was verified using Vinculin, or GAPDH, or α-Tubulin as a loading control, depending on the molecular weight range available on each membrane to avoid overlap with target proteins. Band intensities were quantified by densitometry using ImageJ software and normalized to the corresponding loading control signal.

### Proteomics

Quantitative proteomic analyses were performed on whole-cell lysates obtained from RPE1 WT and PFN1^KO^ cells. Samples were analyzed in quadruplicate. Cells were cultured in Petri dishes and harvested at sub-confluent conditions to avoid confounding effects of contact inhibition. Protein extracts were prepared using RIPA buffer. Proteins were precipitated with ice-cold acetone and resuspended. Protein reduction was carried out in 25 µL of 100 mM NH₄HCO₃ by adding 2.5 µL of 200 mM DTT (Merck) and incubating at 60 °C for 45 min. Samples were subsequently alkylated with 10 µL of 200 mM iodoacetamide (Merck) for 1 h at room temperature in the dark. Excess iodoacetamide was quenched by adding 200 mM DTT. Proteins were then digested with trypsin. The resulting peptide mixtures were dried in a SpeedVac and desalted. Digested peptides were analyzed using an Ultimate 3000 RSLC nano system coupled online to an Orbitrap Exploris 480 equipped with a High-Field Asymmetric Waveform Ion Mobility Spectrometry (FAIMS) interface (Thermo Fisher Scientific). Samples were loaded onto a reversed-phase C18 column (15 cm × 75 µm i.d., Thermo Fisher Scientific) and separated using a 41 min gradient from 6% to 95% mobile phase B at 500 nL/min, followed by a 1 min re-equilibration at 6% mobile phase B. MS data were acquired in positive ion mode using a spray voltage of 2500 V, with the FAIMS interface operated in standard resolution at a compensation voltage (CV) of −45 V. Data were collected in data-independent acquisition (DIA) mode with a precursor m/z range of 400–900, an isolation window of 8 m/z with 1 m/z overlap, HCD collision energy of 27%, Orbitrap resolution of 30,000, and RF lens set to 50%. The normalized AGC target was 1000, maximum injection time was 25 ms, and microscan was set to 1. DIA data were processed using DIA-NN (v1.8.1) with library-free search and deep learning-based prediction of spectra, retention times (RTs), and ion mobility (IMs) enabled. Trypsin/P was specified as the protease; precursor charge states 1–4, peptide lengths of 7–30 amino acids, and precursor m/z values between 400 and 900 were considered, allowing up to two missed cleavages. Carbamidomethylation of cysteine was set as a fixed modification and methionine oxidation as a variable modification (maximum two variable modifications per peptide). The false discovery rate (FDR) was controlled at 1%.

### Cell cycle phase annotation of the proteomic dataset

Cell cycle phase–resolved protein annotations were obtained from Supplementary Table 2 of Rega *et al* ^44^. In the Proteome sheet, proteins were filtered based on the Cell Cycle Dependency column, retaining only entries annotated as ‘CCD’. To define unchanged G1/S- proteins, the “Peak Cell Cycle Phase column” was filtered for Late_G1, G1/S, S, and Palbo. All resulting G1/S-associated proteins were exported into a dedicated file and matched to our RPE1 proteomic dataset. For each protein, the raw abundance values of the four biological replicates for WT and KO cells were reported, together with the corresponding fold-change (FC) values and *p* values calculated using Welch’s *t*-test. For each KO clone, unchanged G1/S proteins were defined as proteins showing FC > -0.7 and < 1.5 (corresponding to log2FC > -0.6 and < 0.6) and *p* > 0.05. The lists of unchanged proteins obtained for each clone were then compared, and only proteins shared between KO#5 and KO#33 were retained.

To define unchanged mitotic proteins, the “Peak Cell Cycle Phase” column was filtered for G2/M_1 and M/Early G1, again considering only CCD proteins. These proteins were matched to our RPE1 proteomic dataset, and the corresponding raw abundance values for the four biological replicates of WT and KO cells were collected, together with relative fold changes and *p* values from Welch’s *t*-test. Unchanged mitotic proteins were defined as proteins with FC > -0.7 and < 1.5 in each KO clone. The overlap between the unchanged protein sets from KO#5 and KO#33 was then determined, yielding 49 shared proteins.

### Mitotic Engagement Score (MES) calculation

To quantitatively assess the degree of mitotic program activation, we computed a Mitotic Engagement Score (MES) based on the relative abundance of selected mitotic regulators representing key functional steps of mitotic progression. Four proteins were chosen to capture sequential stages of mitosis: CDK1, representing the commitment to mitotic entry at the G2/M transition; PRC1, required for spindle midzone formation and proper spindle organization; NDC80, a core component of the kinetochore complex mediating chromosome attachment to spindle microtubules during metaphase; and UBE2C, an E2 ubiquitin-conjugating enzyme essential for APC/C activation, enabling the metaphase-to-anaphase transition and chromosome segregation. For each sample, normalized protein abundance values obtained from quantitative proteomics were used. To control for global variations in mitotic protein expression and ensure comparability across conditions, values were normalized to AURKA, which displayed stable expression across experimental groups and served as an internal mitotic reference. The MES was calculated as the mean normalized abundance of CDK1, PRC1, NDC80, and UBE2C relative to AURKA for each biological replicate.

### In vivo experiments and histological analysis

All animal experiments were conducted in accordance with Italian regulations for animal experimentation and were approved by the Institutional Animal Care and Use Committee (IACUC) (approval number: 1012/2024-PR). Six-week-old male NOD scid gamma (NSG) mice were purchased from Charles River Laboratories and housed under controlled environmental conditions with ad libitum access to food and water.

For intra-tibial xenograft experiments, SaOS2 cells were prepared as single-cell suspensions and 1 × 10⁶ cells were injected into the right tibia of each mouse in a total volume of 10 µL of sterile phosphate-buffered saline (PBS). All procedures were performed under sterile conditions. Mice were anesthetized by intraperitoneal injection of ketamine (100 mg/kg) and xylazine (10 mg/kg), and cell injections were performed using a 26-gauge needle. Mice were monitored weekly for general health status and tumor development. Starting from week 4 post-injection, tumor growth was assessed by caliper measurements. Six weeks after cell injection, mice were euthanized by carbon dioxide inhalation. Following sacrifice, soft tissues and bones were dissected and fixed in 4% paraformaldehyde in PBS (pH 7.4) for 48 h. Soft tissues were subsequently dehydrated through a graded ethanol series (70%, 80%, 90%, and 100%) and paraffin-embedded. For lung metastasis assessment, serial 5-µm-thick lung sections were collected and stained with hematoxylin and eosin (H&E). Histological analysis was performed on sections encompassing the entire tumor area represented in each section. For H&E staining, sections were deparaffinized in xylene for 30 min and rehydrated through a graded ethanol series (100%, 90%, 80%, 70%, and tap water, 5 min each). Sections were stained with hematoxylin solution (Sigma-Aldrich) for 3 min, rinsed in distilled water, and counterstained with 1% eosin solution (Carlo Erba Reagents) for 30 seconds. After dehydration through graded ethanol solutions, sections were mounted with coverslips and analyzed by bright-field microscopy using a Nikon Intensilight C-HGFI system.

### Statistics and Reproducibility

Unless otherwise stated, quantitative data were obtained from two or three independent biological experiments. Statistical analyses were performed using GraphPad Prism software (version 10.6.1 for Mac). The specific statistical tests applied for each experiment are indicated in the corresponding figure legends. Representative images shown in the figures were selected from datasets generated from at least two independent experiments.

## Acknowledgements

Authors acknowledge members of the “Bone Diseases and Tumors” laboratory at IGB-CNR and members of the “Biology of centrosomes and genetic instability” laboratory at Institut Curie for constructive feedback provided during manuscript preparation.

We are grateful to members of the Integrated Microscopy and FACS Facilities of IGB-CNR. We thank the Cell and Tissue Imaging (PICT-IBiSA), member of the French National Research Infrastructure France-BioImaging (ANR10-INBS-04), and the Nikon Imaging center from Institut Curie for microscopy.

We thank Dr Michela Rossi (Bambino Gesù Children’s Hospital, Rome, Italy) for technical assistance with the intratibial injections in NSG mice. We are grateful to Dr Gennaro Gambardella (Telethon Institute of Genetics and Medicine, Naples, Italy), for bioinformatic support in the analysis of the correlation between whole-genome doubling (WGD) and *PFN1* loss across tumor datasets. We thank Dr. Marco Corona (IGB-CNR) for his expert veterinary support and for performing anesthesia procedures during NSG mouse xenograft generation experiments.

F.S.d.C. was supported by AIRC Fellowship for Abroad and LabEx Cell(n)Scale Post-doctoral Fellowship.

The research leading to these results has received funding from the Italian Association for Cancer Research (AIRC; project ID 25110) to F.G., by the Ministry of University and Research under the PRIN 2022 PNRR Call (project code P20224JCNN) to F.G.; and by the Next Generation EU programme within the framework of the National Recovery and Resilience Plan (PNRR), Investment PE8 – Project Age-It, to F.G. M.M. acknowledges funding from AIRC UniCanVax grant n. 22757.

R.B. acknowledges funding from InCA (www.e-cancer.fr) (2021-1- PREV-Bio grant) and the government through the Agence Nationale de la Recherche from France 2030 (ANR-23-CHBS-0012).

## Author contributions

F.S.d.C. and F.G. conceived and designed the study. F.S.d.C. performed the majority of the experiments and analyzed the data. Live-cell imaging and immunofluorescence experiments were performed by F.S.d.C. at the Institut Curie under the supervision of R.B. S.R. contributed to flow cytometry analysis, xenograft generation and histological analyses. S.G. provided conceptual input and methodological guidance throughout the study. N.V. and M.A.B. contributed to cell-based experiments and data acquisition. A.- S.M. developed the Fiji plugin used for the quantitative analysis of FUCCI live-cell imaging and provided technical support for its implementation. M.M. performed and analyzed the proteomic experiment. I.H. and F.F. performed single-cell whole-genome sequencing and conducted the associated bioinformatic analyses. F.S.d.C. wrote the manuscript, with input from F.G., R.B., and S.G. All authors reviewed and approved the final manuscript.

## Data availability

The TCGA Pan-Cancer Atlas data used in this study are publicly available through the Genomic Data Commons PanCanAtlas resource. The osteosarcoma/Paget’s disease of bone whole-genome sequencing data analyzed in this study were previously published and are available as described in Scotto di Carlo et al, 2023 (ref.^15^).

The mass spectrometry proteomics data and single-cell whole-genome sequencing data generated in this study will be deposited in appropriate public repositories before publication. Source data underlying the main and supplementary figures are provided with this paper as Supplementary Data files. Any additional information required to reanalyze the data reported in this paper is available from the corresponding authors upon reasonable request.

## Code availability

Custom code used for the TCGA copy-number and whole-genome doubling analyses, together with the custom Fiji/ImageJ plugin used for quantitative analysis of FUCCI live-cell imaging, will be made available from the corresponding authors upon reasonable request. No other custom code was used to generate the data reported in this study.

