## Supplementary Figures for "Profilin 1 maintains cell cycle fidelity to prevent unscheduled genome doubling and polyploidy in cancer"

[a]Institute of Genetics and Biophysics "Adriano Buzzati-Traverso", National Research Council of Italy, Naples, Italy.
[b]Biology of centrosomes and genetic instability, Institut Curie, PSL Research University, CNRS UMR 144, Paris, France.
[c]Cell and Tissue Imaging Facility (PICT-IBiSA), Institut Curie, CNRS UMR144 PSL Research University, Paris, France
[d]European Research Institute for the Biology of Ageing, University of Groningen, University Medical Center Groningen, Groningen, The Netherlands.
[e]Department of Translational Medicine, Università del Piemonte Orientale, 28100 Novara, Italy.
[f]Institute for Molecular and Translational Cardiology (IMTC), IRCCS Policlinico San Donato, S. Donato Milanese, 20097, Italy

This file contains 7 Supplementary Figures and Legends.

### Supplementary Figure 1

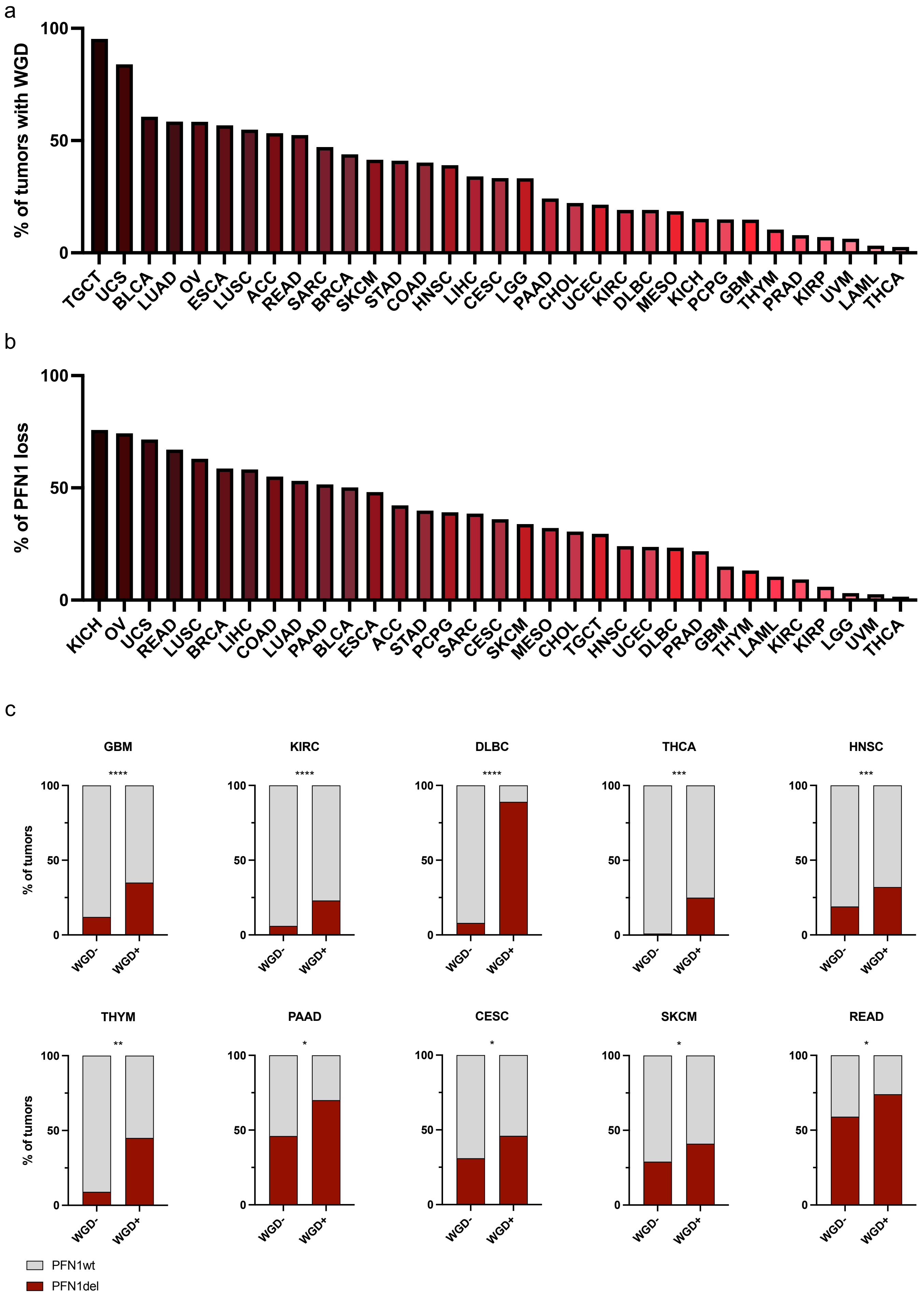

**Supplementary Figure 1**:
**a.** Frequency (%) of tumors exhibiting whole-genome doubling (WGD) across cancer types in the TCGA Pan-Cancer Atlas. Tumor abbreviations and sample numbers are provided in **Supplementary Data 1**.
**b.** Frequency (%) of tumors harboring loss of at least one *PFN1* copy across cancer types in the TCGA Pan-Cancer Atlas. Tumor abbreviations and sample numbers are provided in **Supplementary Data 1.**
**c.** Stacked bar plots showing the distribution of tumors with or without whole-genome doubling (WGD), stratified by PFN1 status (WT or DEL), across different cancer types. Tumor abbreviations and sample numbers are provided in **Supplementary Data 1.** Data are shown as the percentage of tumors within each cohort. Asterisks indicate statistical significance assessed by two-sided Fisher's exact test. *P*-values were calculated using absolute tumor counts, whereas percentages are shown for visualization. *P*-values are as follows: GBM, $p < 0.0001$; KIRC, $p < 0.0001$; DLBC, $p < 0.0001$; THCA, $p = 0.0004$; HNSC, $p = 0.0007$; THYM, $p = 0.0057$; PAAD, $p = 0.0104$; CESC, $p = 0.0146$; SKCM, $p = 0.0171$; READ, $p = 0.0458$. Corresponding odds ratios (OR), 95% confidence intervals (CI), and false discovery rate (FDR)–adjusted *p*-values are reported in **Table 1**.

#### Supplementary Figure 2

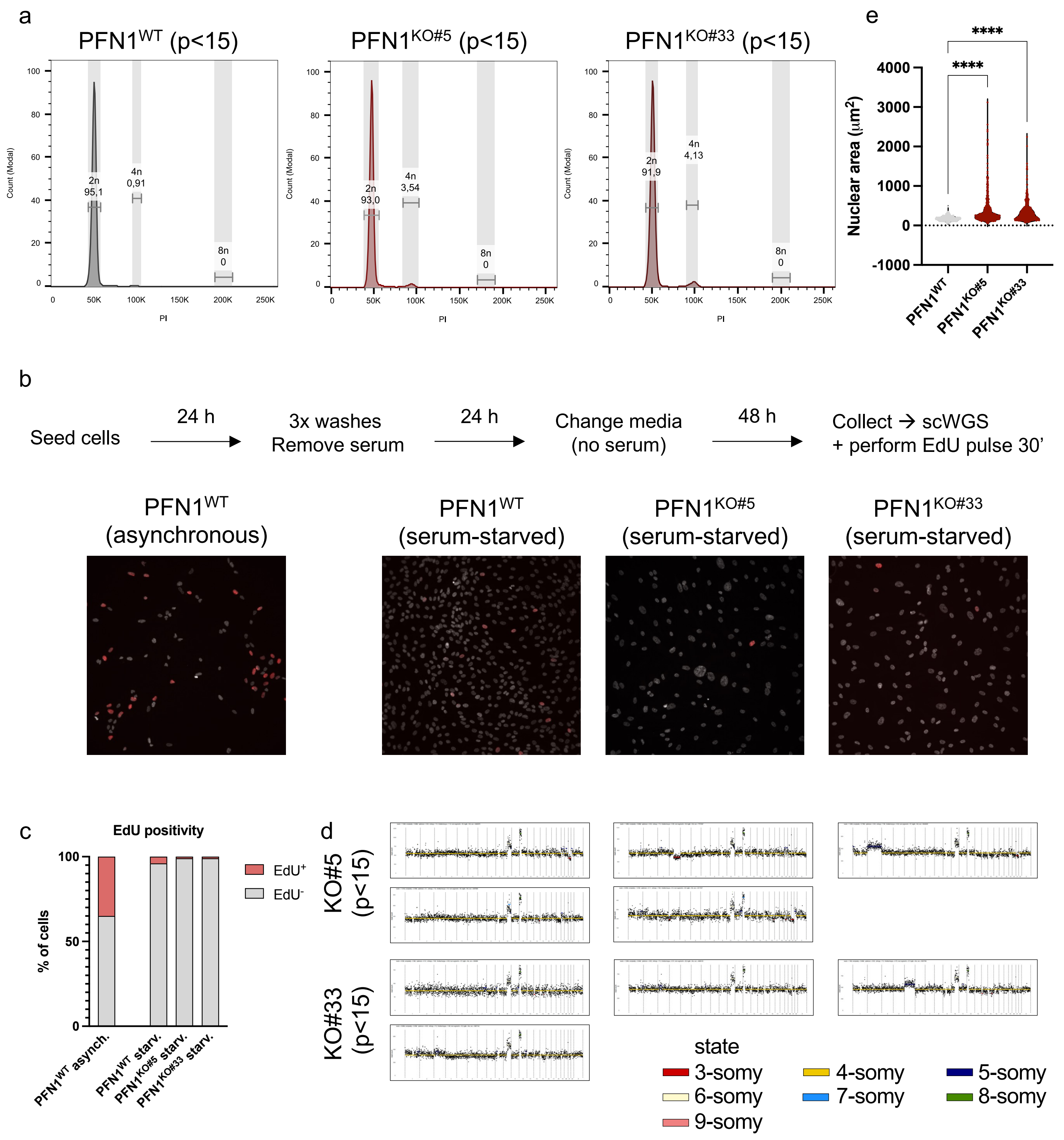

**Supplementary Figure 2**:

**a.** DNA content analysis by propidium iodide (PI) staining in serum-starved early-passage (p<15) WT and $PFN1^{KO}$ (KO#5 and KO#33) RPE1 cells. Histograms show cell counts expressed in modal values plotted against DNA content measured by PI incorporation. Diploid (2n) and tetraploid (4n) peaks are highlighted and relative abundance (%) is shown.

**b.** Schematic overview of the serum-starvation protocol. Cells were serum-starved for 72 h; an aliquot of the same cell populations processed for single-cell whole-genome sequencing (scWGS) was pulsed with EdU for 30 minutes to verify effective cell cycle arrest prior to sequencing. Representative images below show EdU incorporation (red) in asynchronous WT RPE1 cells (positive control) and in serum-starved WT and $PFN1^{KO}$ cells. Nuclei are counterstained with DAPI (grey).

**c.** Stacked bar plot showing the quantification of EdU-positive ($EdU^{+}$) and EdU-negative ($EdU^{-}$) cells, expressed as percentage of total cells analyzed. *N* = 425 cells for asynchronous WT; 2395 cells for WT; 1004 cells for KO#5; 902 cells for KO#33.

**d.** Single-cell whole-genome copy-number profiles from tetraploid (4n) $PFN1^{KO\#5}$, $PFN1^{KO\#10}$ and $PFN1^{KO\#33}$ RPE1 cells following serum starvation. Data points are color-coded according to inferred ploidy states; different ploidy states are indicated by distinct colors.

**e.** Quantification of nuclear area measured at the single-cell level in $PFN1^{WT}$ and $PFN1^{KO}$ RPE1 cells (clone #5 and clone #33). Each dot represents one nucleus; *n* = 635 nuclei for WT; 492 nuclei for KO#5; 532 nuclei for KO#33. Statistical significance was assessed using ordinary one-way ANOVA; adjusted *p*-value < 0.0001. Nuclear area is expressed in $\mu m^2$.

### Supplementary Figure 3

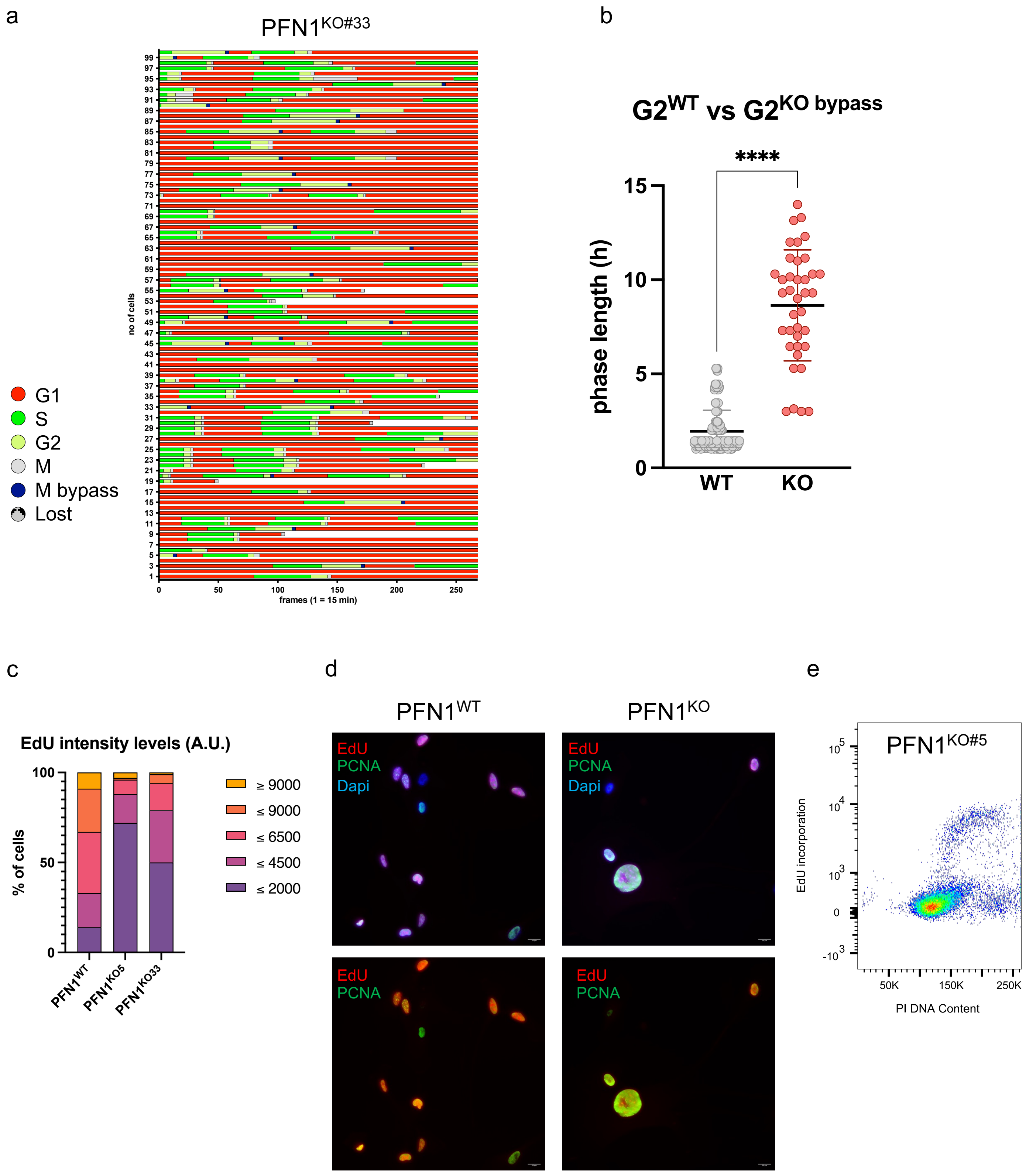

**Supplementary Figure 3**:

**a.** Duration and sequence of cell cycle phases $PFN1^{KO\#33}$ RPE1 cells, as determined by FUCCI-tracking. Each horizontal lane represents a single cell (n = 100 cells per condition). The x-axis represents time in imaging frames (1 frame = 15 minutes). Cell cycle phases are color-coded as follows: G1 (red), S (green), G2 (yellow), mitosis (grey), mitotic bypass (blue), and lost cells (grey with black dots).

**b.** Dot plots showing the duration (in hours) of individual cells in G2 in $PFN1^{WT}$ (grey) and $PFN1^{KO}$ (pink with red contour) RPE1 cells. For $PFN1^{KO}$, only G2 phases from cells that subsequently underwent mitotic bypass are shown. Data from KO#5 and KO#33 clones were pooled. The number of analyzed cells was as follows: WT, $n$ = 177; KO, $n$ = 38. Statistical significance was assessed using Welch's t-test; $p < 0.0001$.

**c.** Stacked bar plot showing the distribution of EdU fluorescence intensity levels in $PFN1^{WT}$, $PFN1^{KO\#5}$, and $PFN1^{KO\#33}$ RPE1 cells. Cells were stratified into intensity categories as indicated (≤2000, ≤4500, ≤6500, ≤9000, and ≥9000 arbitrary units, A.U.) and expressed as percentage of total cells per condition. The number of analyzed cells was: $n$ = 138 (WT), 146 (KO#5), and 131 (KO#33).

**d.** Representative images of $EdU^{+}$ cells (red) co-stained for PCNA (green). EdU labeling was performed for 18 h, and PCNA staining was used to support ongoing DNA synthesis in cells with enlarged nuclei. Nuclei are counterstained with DAPI (blue). Scale bar, 20 µm.

**e.** Flow cytometry analysis of EdU incorporation combined with DNA content measurements in late-passage $PFN1^{KO\#5}$ RPE1 cells. The plot displays EdU fluorescence versus propidium iodide (PI) signal, enabling the visualization of DNA synthesis in cells with different DNA contents.

#### Supplementary Figure 4

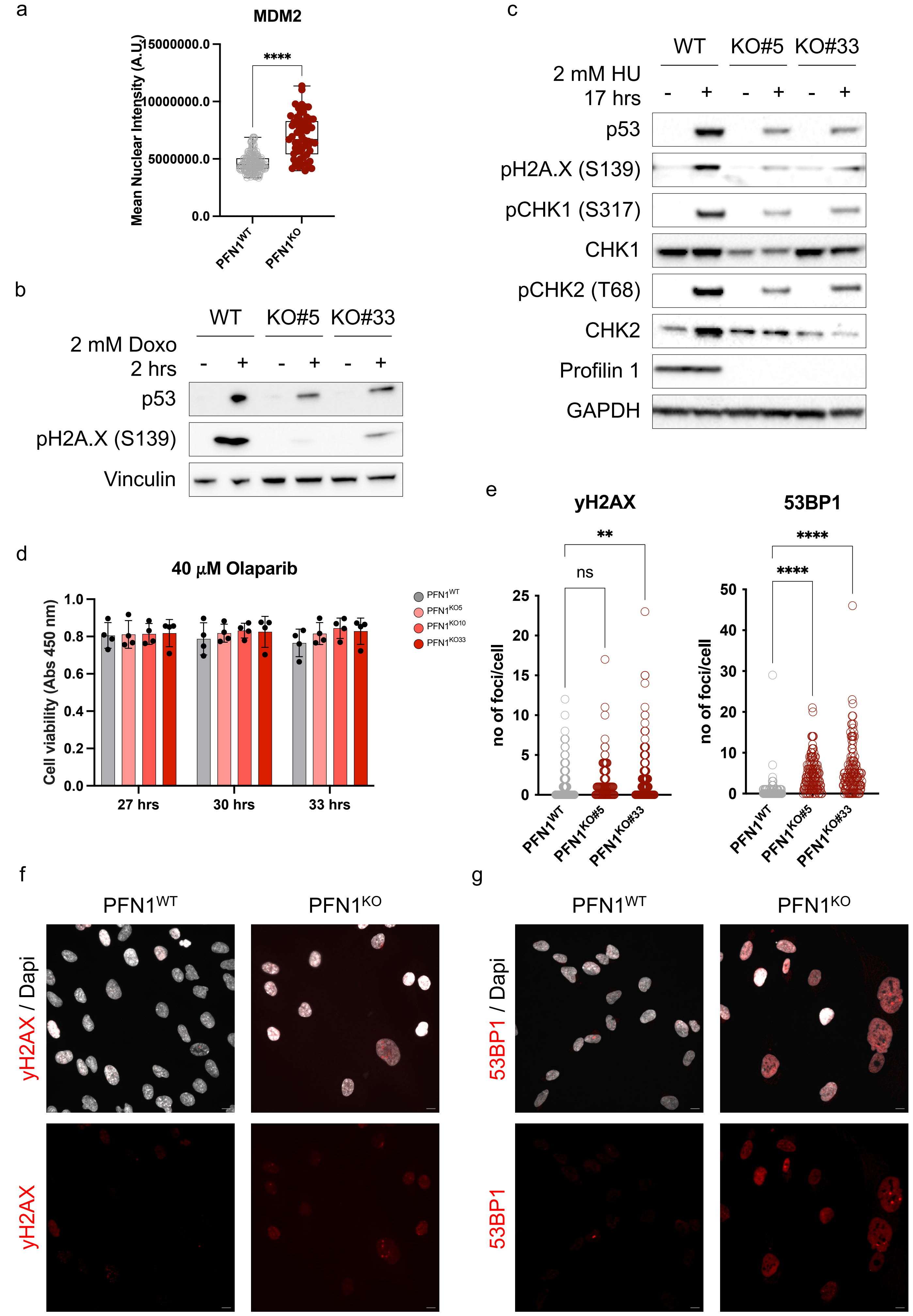

**Supplementary Figure 4**:

**a.** Box plot showing the mean fluorescence intensity of nuclear MDM2 in WT and $PFN1^{KO}$ RPE1 cells. Statistical analysis was performed through unpaired t-test; ****$p < 0.0001$; $n$ = 242 nuclei for WT; 61 for KO#33.

**b.** Western blot analysis of p53 expression in WT and $PFN1^{KO}$ cells treated or not with doxorubicin (Doxo; 2 µM) for 2 hours. Phosphorylation of H2A.X (γH2AX) was used as readout of the DNA damage response; Vinculin was used as a loading control.

**c.** Western blot analysis of p53 expression in WT and $PFN1^{KO}$ cells treated or not with hydroxyurea (HU; 2 mM) overnight. Phosphorylation of H2A.X (γH2AX), CHK1, and CHK2 were used as readouts of the DNA damage response; GAPDH was used as a loading control.

**d.** Cell viability assays (CCK-8) showing absorbance at 450 nm for WT and $PFN1^{KO}$ cells following exposure to 40 µM Olaparib for the indicated durations. Statistical analyses were performed using 2way ANOVA; all multiple comparisons resulted non-significant ($p > 0.05$).

**e.** Quantification of the number of γH2AX (left) and 53BP1 (right) foci per cell in WT and $PFN1^{KO}$ cells. Statistical analyses were performed using ordinary one-way ANOVA. For γH2AX, ns $p$ = 0.3282, **$p$ = 0.0057; $n$ = 185 cells for WT, 102 for KO#5, 148 for KO#33. For 53BP1, ****$p < 0.0001$; $n$ = 190 cells for WT, 126 for KO#5, 127 for KO#33.

**f.** Representative immunofluorescence images of WT and $PFN1^{KO}$ RPE1 cells stained for γH2AX (red) under basal, untreated conditions. Nuclei are counterstained with DAPI (grey). Scale bar, 20 µm.

**g.** Representative immunofluorescence images of WT and $PFN1^{KO}$ RPE1 cells stained for 53BP1 (red) under basal, untreated conditions. Scale bar, 20 µm.

### Supplementary Figure 5

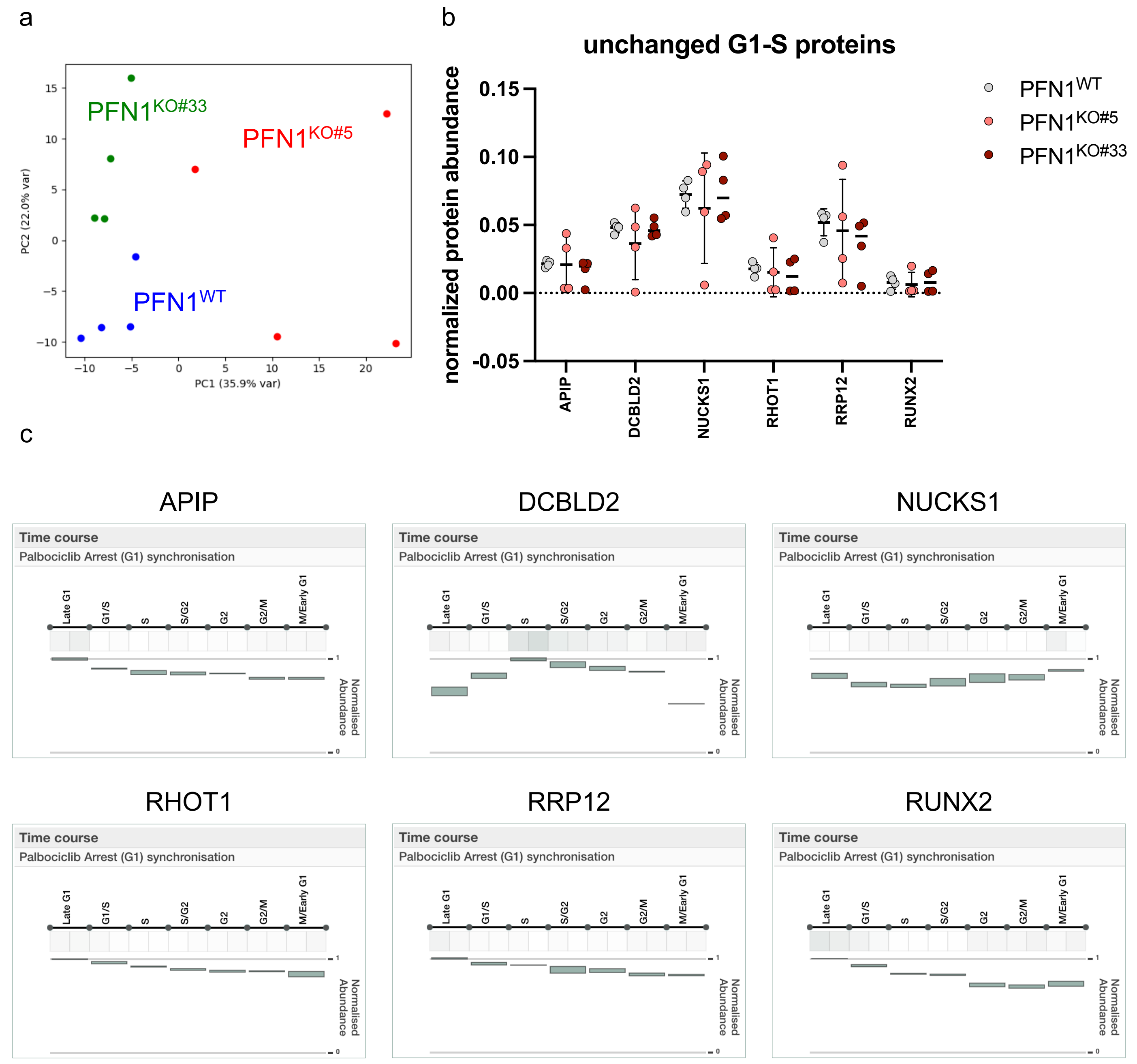

**Supplementary Figure 5**:

**a.** Principal component analysis (PCA) of proteomics samples showing clustering of WT and $PFN1^{KO}$ RPE1 cells based on global protein abundance profiles. Each point represents one biological replicate.

**b.** Graph showing normalized abundance of the indicated proteins derived from proteomic analysis of WT and PFN1KO RPE1 cells. Each dot of the same color represents an independent replicate of the corresponding clone. Statistical analysis was performed using an unpaired Welch's t-test; all comparisons were non-significant ($p > 0.05$).

**c.** Temporal expression profiles of the indicated proteins across the cell cycle, retrieved from the publicly available SLiM cell cycle proteomics database (https://slim-tools.org/cell_cycle/), based on quantitative proteomics of cells synchronized in G1 by Palbociclib treatment and sampled across successive cell cycle stages, as described in the reference **43** (Rega, C. et al. High resolution profiling of cell cycle-dependent protein and phosphorylation abundance changes in non-transformed cells. Nat. Commun. 16, (2025)). Proteins shown were selected based on their peak abundance during G1 and S phases.

**Supplementary Figure 6**

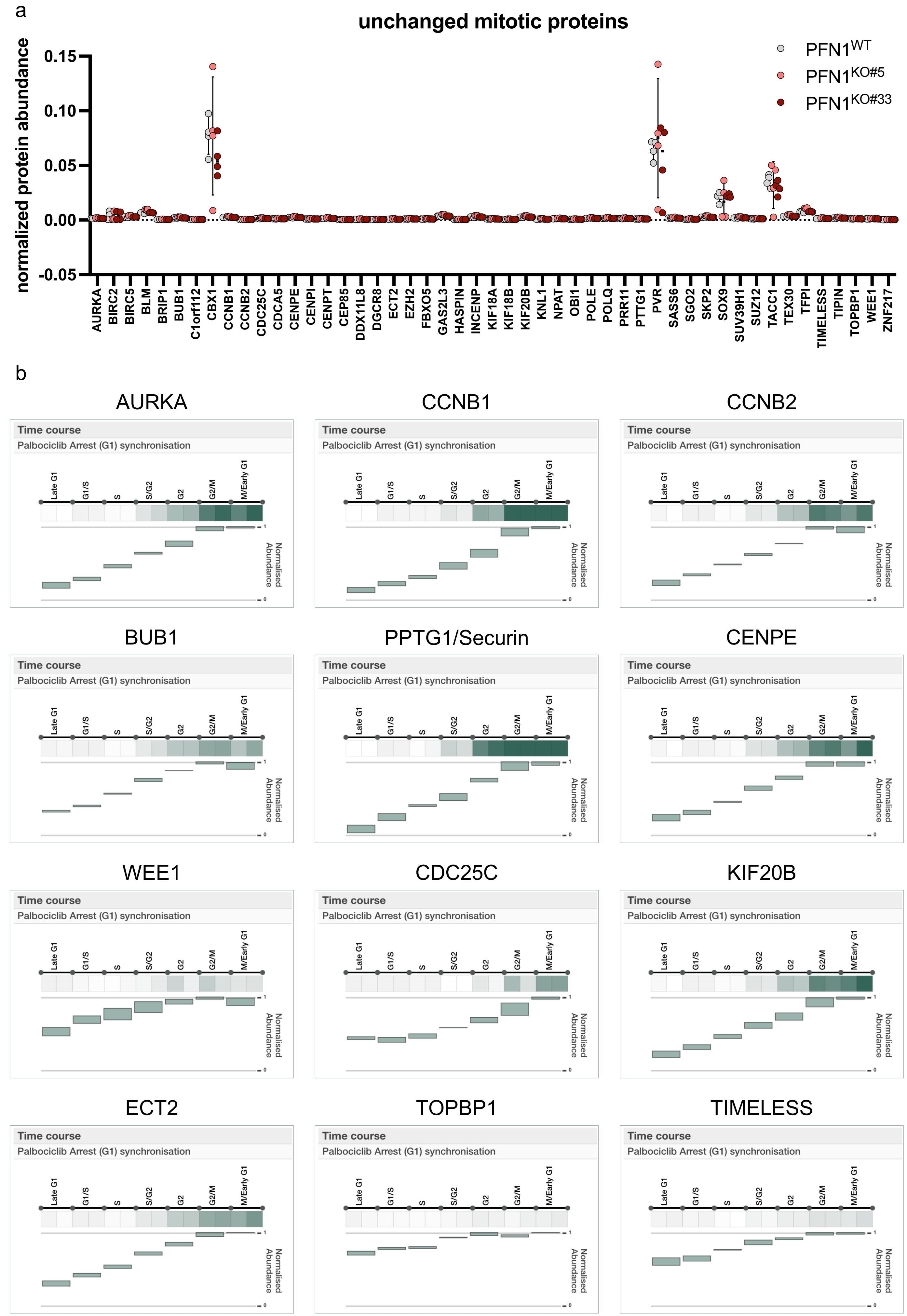

**Supplementary Figure 6**:

**a.** Graph showing normalized abundance of the indicated proteins derived from proteomic analysis of WT and PFN1$^{KO}$ RPE1 cells. Each dot of the same color represents an independent replicate of the corresponding clone. Statistical analysis was performed using an unpaired Welch's t-test; all comparisons were non-significant ($p > 0.05$).

**b.** Temporal expression profiles of the indicated proteins across the cell cycle, retrieved from the publicly available SLiM cell cycle proteomics database (https://slim-tools.org/cell_cycle/), based on quantitative proteomics of cells synchronized in G1 by Palbociclib treatment and sampled across successive cell cycle stages, as described in the reference **43** (Rega, C. et al. High resolution profiling of cell cycle-dependent protein and phosphorylation abundance changes in non-transformed cells. Nat. Commun. 16, (2025)). Proteins shown were selected based on their peak abundance during G2/M and M phases.

**Supplementary Figure 7**

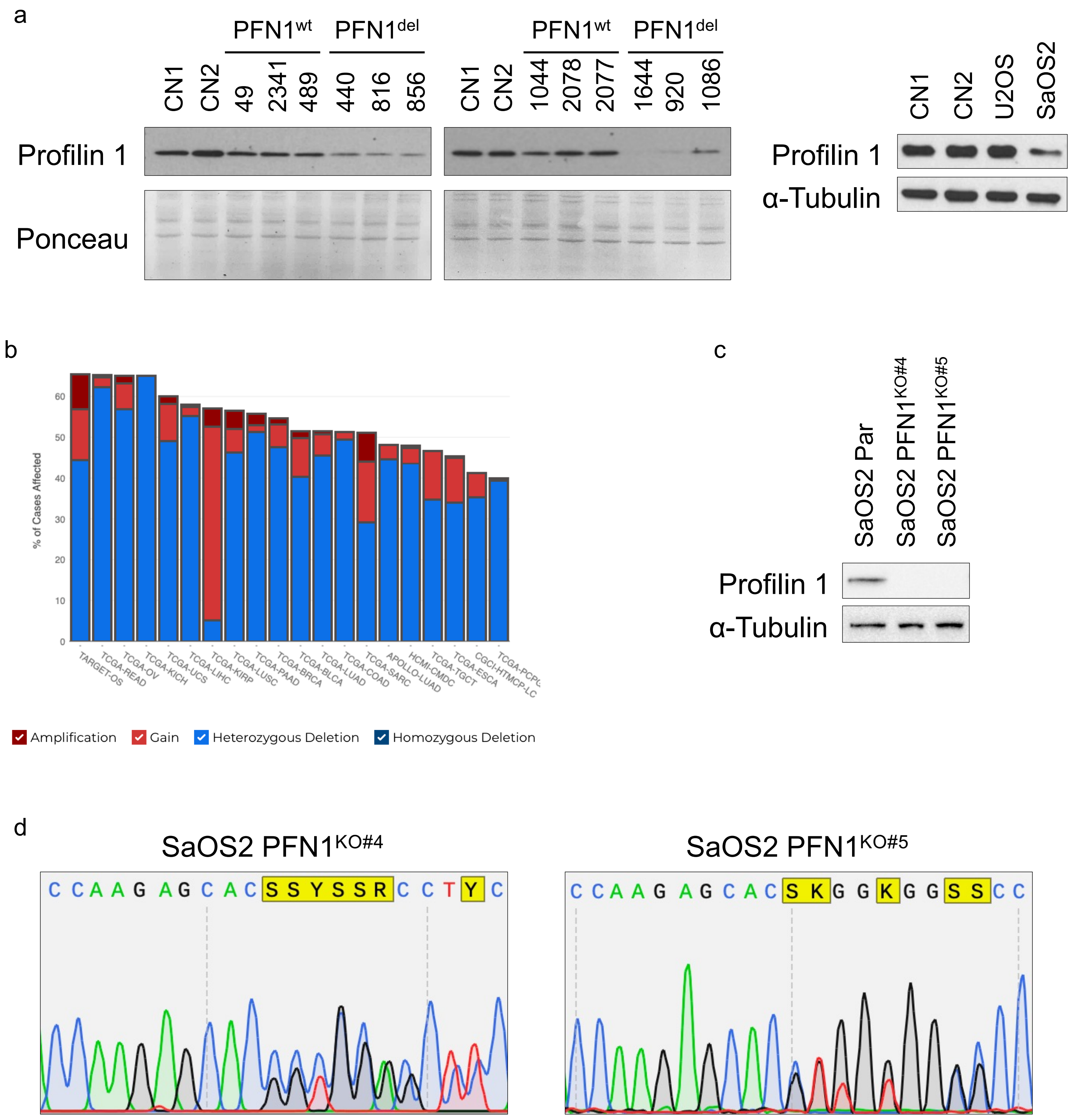

**Supplementary Figure 7**:

**a.** Western blot analysis of Profilin 1 expression in 12 primary osteosarcoma (OS) samples compared with two non-tumoral mesenchymal cell lines (CN1 and CN2). Samples were run on two separate gels. $PFN1^{WT}$ indicates samples in which both *PFN1* gene copies were detected; $PFN1^{DEL}$ indicates OS samples in which loss of heterozygosity at the *PFN1* locus was detected. Ponceau staining was used as a loading control. On the right, western blot analysis of Profilin 1 expression in two OS cell lines (U2OS and SaOS2) compared with two non-tumoral mesenchymal cell lines (CN1 and CN2). α-Tubulin was used as a loading control.

**b.** TCGA-derived data showing genomic alterations affecting the *PFN1* locus across the indicated tumor types, including amplification, gain, heterozygous deletion, and homozygous deletion.

**c.** Western blot analysis of Profilin 1 expression in parental SaOS2 cells (Par) and two independent CRISPR/Cas9-engineered PFN1 knockout SaOS2 clones (#4 and #5). α-Tubulin was used as a loading control.

**d.** Representative Sanger sequencing electropherograms showing CRISPR/Cas9-induced insertions and deletions (indels) at the PFN1 genomic locus in two independent SaOS2 knockout clones.
